# Evidence suggesting creatine as a new central neurotransmitter: presence in synaptic vesicles, release upon stimulation, effects on cortical neurons and uptake into synaptosomes and synaptic vesicles

**DOI:** 10.1101/2022.12.22.521565

**Authors:** Xiling Bian, Jiemin Zhu, Xiaobo Jia, Wenjun Liang, Sihan Yu, Zhiqiang Li, Wenxia Zhang, Yi Rao

## Abstract

The discovery of a new neurotransmitter, especially one in the central nervous system (CNS), is both important and difficult. We have been searching for new neurotransmitters for 12 years. We detected creatine (Cr) in synaptic vesicles (SVs), at a level lower than glutamate (Glu) and gamma-aminobutyric acid (GABA) but higher than acetylcholine (ACh) and 5-hydroxytryptamine (5-HT). SV Cr was reduced in mice lacking either arginine:glycine amidinotransferase (AGAT, a Cr synthetase) or SLC6A8, a Cr transporter with mutations among the most common causes of intellectual disability (ID) in men. Calcium-dependent release of Cr was detected after stimulation in brain slices. Cr release was reduced in SLC6A8 and AGAT mutants. Cr inhibited neocortical pyramidal neurons. SLC6A8 was necessary for Cr uptake into synaptosomes. Cr was found by us to be taken up into SVs in an ATP dependent manner. Our biochemical, chemical, genetic and electrophysiological results are consistent with the possibility of Cr as a neurotransmitter, though not yet reaching the level of proof for the now classic transmitters. Our novel approach to discover neurotransmitters is to begin with analysis of contents in SVs before defining their function and physiology.

## Introduction

Neural signaling depends on chemical transmission between neurons and their target cells ^1–6^. Neurotransmission depends on chemicals such as neurotransmitters, neuromodulators and neuropeptides. Decades of work, sometimes with convoluted paths, were involved before a molecule was established as a classic neurotransmitter ^7^. Initial hints about cholinergic signaling were obtained in the 1800s ^8–11^. Choline ^12,13^ and acetylcholine (ACh) ^14^ were discovered decades before their pharmacological effects were found around the turn of 20^th^ century ^15–17^. Henry Dale and colleagues realized similarities of ACh and parasympathetic stimulation ^5,18^ but it was not until 1929 when ACh was detected in the body ^19^ and 1934 when ACh was proven a neurotransmitter in the peripheral nervous system (PNS) ^20–22^. It took nearly 100 years from the finding of the effects of supradrenal gland damage ^23^ or removal ^24^, the observation of an activity in the supradrenal gland ^25^, the isolation of an inactive derivative ^26–29^, and the successful isolation of adrenaline ^30–32^, the notice of similarities between adrenaline and sympathetic stimulation ^3,4,33–35^, to the mid-1940s when Ulf von Euler proved that noradrenaline (NA) was the neurotransmitter of the sympathetic nerves ^36–38^. While it is not easy to establish a molecule as a neurotransmitter in the PNS, it is even harder to establish a central nervous system (CNS) neurotransmitter. Three decades elapsed between the time when ACh was proven to be a PNS neurotransmitter and the time when it was established as a CNS neurotransmitter ^39–41^ and two decades between NA as a peripheral transmitter and a central transmitter ^42,43^.

If a neurotransmitter acts only in the CNS, but not in the PNS, it is much more difficult to discover or to prove. Most neurotransmitters were discovered for their effects on peripheral tissues, with muscle contraction or relaxation as a major readout. Glutamate (Glu) ^44–47^ and gamma-aminobutyric acid (GABA) ^46,48–51^ were discovered partly because of their peripheral effects and partly because of their effects on spinal neurons. There is no reason for central neurotransmitters to also act peripherally, but relatively little efforts have been reported to find small molecule neurotransmitters acting only on CNS neurons with no peripheral bioassays available. Premature assumptions and technical difficulties are among the major reasons why the hunt for neurotransmitters has not been a highly active area of research over the last three decades.

Are there more neurotransmitters and how can they be discovered? Classic neurotransmitters are stored in synaptic vesicles (SVs) ^52–60^. They are released upon electric stimulation before being degraded enzymatically or taken up into the presynaptic terminal by cytoplasmic transporters and into SVs by vesicular transporters ^61–66^. Most of the major textbooks list either 3 ^67–69^ or 4 ^70–72^ criteria of a neurotransmitter: presence in presynaptic neurons, release upon stimulation, action on postsynaptic neurons, mechanism of removal. Some molecules commonly accepted as neurotransmitters still do not meet all the criteria listed in different textbooks, but they nonetheless play important functional roles in the CNS and their defects cause human diseases. Over time, different small molecules have been proposed to function as neurotransmitters (e.g., 46,73), but none satisfies all the criteria. Robust and reliable detection of the candidate molecule in SVs is often, though not always, the problem (cf. 74).

Beginning in 2011, we have been actively searching for new neurotransmitters in the mammalian brain. We have tried different approaches, including searching for neuroactive substances in the cerebral spinal fluid (CSF) and following transporters potentially localized in the SVs. One approach that we have now taken to fruition is the purification of the SVs from mouse brains coupled with chemical analysis of their contents. We have found known transmitters such as Glu, GABA, ACh, and 5-hydroxytryptamine (5-HT). But more importantly, we have reproducibly detected creatine (Cr) in SVs.

Cr was discovered in 1832 by Michel-Eugène Chevreul ^75,76^ and has long been considered as an energy buffer in the muscle and the brain ^77–79^. Half of Cr in a mammalian animal is thought to come from diet and the rest from endogenous synthesis ^80^. Most of the Cr is present in the muscle but it is also present in the brain. Although most of the endogenous Cr is synthesized in the kidney, the pancreas and the liver ^77,81^, Cr is also synthesized in the brain ^80,82,83^.

Solute carriers (SLC) contribute to both cytoplasmic and vesicular transporters. With 19 members in humans, family 6 (SLC6) are secondary active transporters relying on electrochemical Na^+^ or H^+^ gradients ^84–89^. SLC6 is also known as the neurotransmitter transporter (NTT) family because some members transport neurotransmitters such as GABA (by SLC6A1 or GABA transporter 1, GAT1; SLC6A13 or GAT2; SLC6A11 or GAT3) ^90–93^, NA (by SLC6A2 or NA transporter,

NET) ^65^, dopamine (by SLC6A3 or DAT) ^94–96^, 5-HT (by SLC6A4 or serotonin transporter, SERT) ^66,97^, and glycine (by SLC6A9, or GlyT1; SLC6A5 or GlyT2) ^90,98,99^. Cr is transported by SLC6A8 (also known as CrT, CT1 or CRCT) ^100–106^. In addition to peripheral organs and tissues, SLC6A8 is also expressed in the nervous system where it is mainly in neurons ^82,104,107–109^. SLC6A8 protein could be found on the plasma membrane of neurons ^109–111^.

The functional significance of SLC6A8 in the brain is supported by symptoms of humans defective in SLC6A8. Mutations in SLC6A8 were found in human patients with intellectual disability (ID), delayed language development, epileptic seizures and autistic-like behaviors ^112,113^. They are collectedly known as Cr transporter deficiency (CTD), with ID as the hallmark. Particular vulnerability of language development has been observed in some SLC6A8 mutations which had mild ID but severe language delay ^114^. CTD contributes to approximately 1 to 2.1% of X-linked mental retardation ^115–123^. While CTD is highly prevalent in ID males, it is also present in females, with an estimated carrier frequency of 0.024% ^124^.

SLC6A8 knockout mice ^125–127^ showed typical symptoms of human CTD patients with early and progressive impairment in learning and memory. Mice with brain- and neuronal-specific knockout of SLC6A8 showed deficits in learning and memory without changes in locomotion caused by peripheral involvement of SLC6A8 ^128,129^. Deletion of SLC6A8 from dopaminergic neurons in the brain caused hyperactivity ^130^. These results demonstrate that SLC6A8 is functionally important in neurons.

Cr deficiency syndromes (CDS) are inborn errors of Cr metabolism, which can result from defects in one of the three genes: guanidinoacetate methyltransferase (GAMT) ^131^, arginine-glycine amidinotransferase (AGAT) ^132,133^, and SLC6A8 ^112^. That they all show brain disorders indicates the functional importance of Cr in the brain^134–137^. Cr uptake was observed previously ^220, 221^.

Here we first biochemically purified SVs from the mouse brain and discovered the presence of Cr, as well as classic neurotransmitters Glu and GABA, ACh and 5-HT, in SVs. We then detected calcium (Ca^2+^) dependent releases of Cr, Glu and GABA but not ACh and 5-HT when neurons were depolarized by increased extracellular concentrations of potassium (K^+^). Both the level of Cr in SVs and that of Cr released upon stimulation were decreased significantly when either the gene for SLC6A8 or the gene for AGAT were eliminated genetically. When Cr was applied to slices from the neocortex, the activities of pyramidal neurons were inhibited. Furthermore, we confirmed that Cr was taken up by synaptosomes and found that Cr uptake was significantly reduced when the SLC6A8 gene was deleted. Finally, we found that Cr was transported into SVs. Thus, multidisciplinary studies with biochemistry, genetics and electrophysiology have suggested that Cr is a new neurotransmitter, though the discovery of a receptor for Cr would prove it.

## Results

### Detection of Cr in SVs from the mouse brain

To search for new neurotransmitters, we tried several approaches. For example, we used Ca^2+^ imaging to detect neuroactive substances in the cerebrospinal fluid (CSF), but it was difficult to rule out existing neurotransmitters and select responses from potentially new neurotransmitters. We also transfected cDNAs for all human SLCs into dissociated cultures of primary neurons from the mouse brain and found that more than 50 out of all SLCs could be localized in SVs. However, when we used CRISPR-CAS9 to tag some of the candidate SLCs in mice, some of them were found to be expressed outside the CNS, indicating that, while ectopic expression of these candidate SLCs could be localized on SVs, the endogenous counterparts were not localized on SVs.

Here we report our approach using the purification of SVs as the first step (Supplementary Figure 1A) ^138–145^. Synaptophysin (Syp) is a specific marker for SVs ^146–149^ and an anti-Syp antibody was used to immunoisolate SVs ^143,144,150–152^. Visualization by electronic microscopy (EM) (Supplementary Figure 1B, left panel) showed that the purified vesicles were homogenous, with an average diameter of 40.44 ± 0.26 nm (n=596 particles) (Supplementary Figure 1B, right panel), consistent with previous descriptions of SVs ^153,154^.

Immunoblot analysis with twenty markers of subcellular organelles of neurons and one marker for glia (Supplementary Figure 1C) indicate that our purifications were highly effective, with SV markers detected after purification with the anti-Syp antibody, but not that with the control immunoglobulin G (IgG). SV proteins included Syp ^146–149^, synaptotagmin (Syt1) ^155^, synatobrevin2 (Syb2) ^156,157^, SV2A ^158^, H^+^-ATPase ^159^, and vesicular neurotransmitter transporters for glutamate (VGLUT1, VGLUT2) ^160^ and GABA (VGAT) ^161^. Immunoisolation by the anti-Syp antibody did not bring down markers for the synaptic membrane (with SNAP23 as a marker) ^162–164^, postsynaptic components (with PSD95 and GluN1 as markers) ^165,166^, the Golgi apparatus (with GM130 and Golgin 97 as markers) ^167,168^, early endosome (with early endosome-associated 1, EEA1, as a marker) ^169^, the lysosome (with LC3B and cathepsinB as markers), the cytoplasma (with glyceraldehyde-3-phophate dehydrogenase or GAPDH as a marker), mitochondria (with voltage-dependent anion channel or VDAC, as a marker), cytoplasmic membrane (with calcium voltage-gated channel subunit alpha 1 or CACNA1A as a marker), axonal membrane (with glucose transporter type 4 or GluT4 as a marker) and glia membrane (with myelin basic protein or MBP as a marker). These results indicated that the SVs we obtained were of high integrity and purity.

To detect and quantify small molecules as candidate transmitters present in the purified SVs, capillary electrophoresis-mass spectrometry (CE-MS) was optimized and utilized (Figure 1A, Supplementary Figure 1A) ^151,170^. We found that the levels of classical neurotransmitters such as Glu, GABA, ACh and 5-HT were significantly higher in SVs pulled down by the anti-Syp antibody than those in lysates pulled down by the control IgG (Figure 1 A-E). Consistent with previous reports ^144,151^, significant enrichment of neurotransmitters was observed only from SVs immunoisolated at near 0 °C, but not at the room temperature (RT) (Figure1 A-E). By contrast, another small molecule, alanine (Figure 1G), was not elevated in SVs as compared to the control.

**Figure 1.**
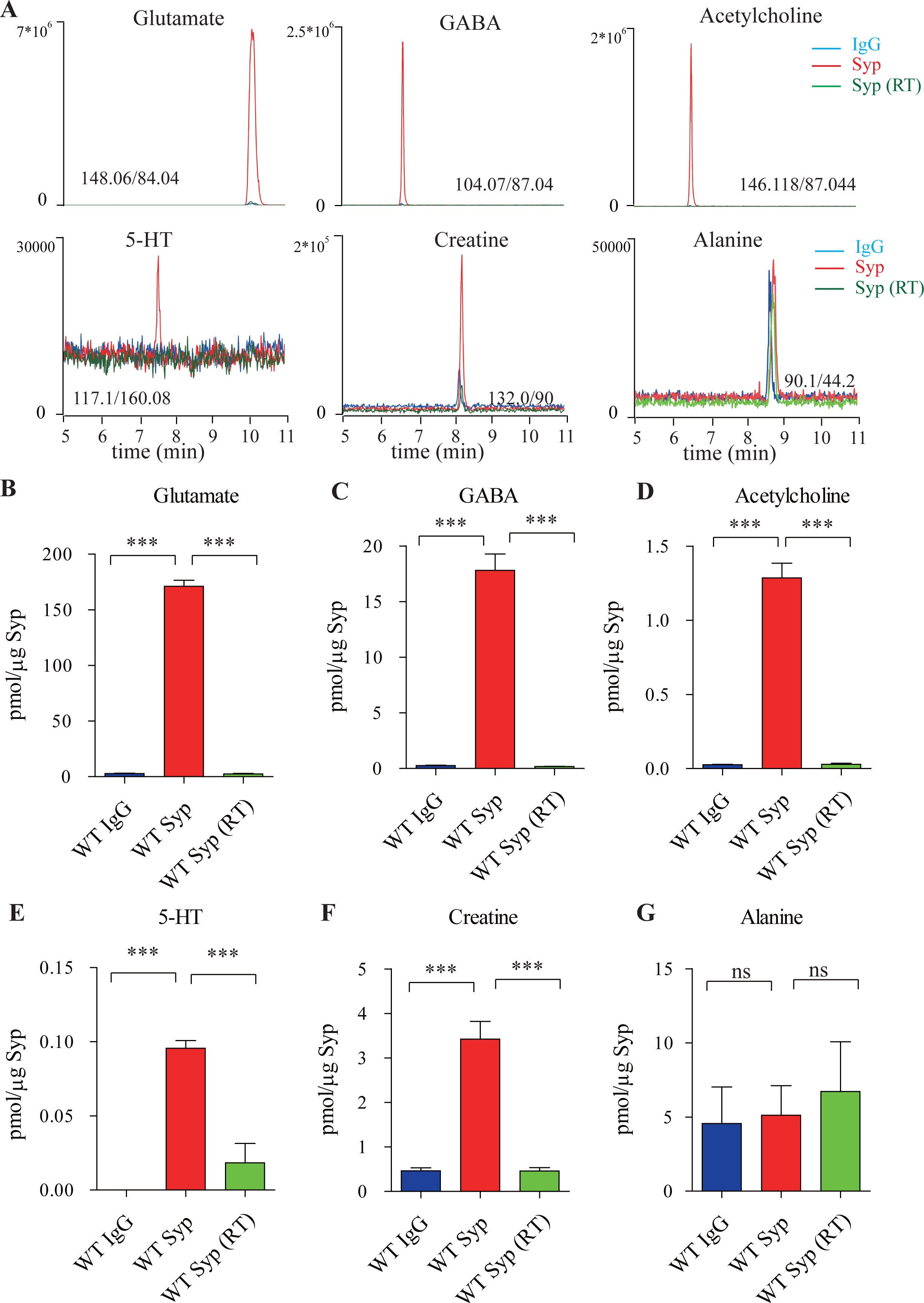
Presence of Cr in SVs from the mouse brain. (A) Representative raw traces from CE-MS of indicated molecules from samples immunoisolated by the control IgG (blue) at 0-2 °C, the monoclonal anti-Syp antibody at 0-2 °C (red) and the anti-Syp antibody at room temperature (RT, green). Q1/Q3 for identifying targets were indicated. (B-G) Quantification of the amounts of indicated molecules. The amount of a molecule was divided by the amount of the anti-Syp antibody bound to magnetic beads. Note Glu (B), GABA(C), ACh (D), 5-HT (E), Cr (F), but not alanine (G) was higher in SVs pulled down by the anti-Syp antibody at 0-2 °C than those pulled down by the IgG control or those pulled down at the RT. n=10 (B-E, and G) or 14 (F) samples per group, ***, p<0.001, ns, not significant. one-way ANOVA with Tukey’s correction.

The amount of Glu was 171.1 ± 5.4 pmol/μg anti-Syp antibody (n=10, Figure 1B), approximately 10 times that of GABA (n=10,17.81 ± 1.47 pmol/μg anti-Syp antibody, Figure 1C). The amount of ACh was 1.29 ± 0.10 pmol/μg anti-Syp antibody (n=10, Figure 1D), approximately 0.072 that of GABA. The amount of 5-HT was 0.096 ± 0.017 pmol/μg anti-Syp antibody (n=10, Figure 1E). Thus, our purification and detection methods were highly reliable and sensitive enough to detect established neurotransmitters.

Under the same conditions, we also detected Cr in SVs (n=14, Figure 1A and 1F). Amount of Cr in the SVs was found to be 3.43 ± 0.40 pmol/μg anti-Syp antibody (Figure 1F), which was approximately 2% of Glu, 19% of GABA, 266% of ACh and 3573% of 5-HT. It is unlikely that these could be attributable to different levels for different neurotransmitters in each SV, but more likely attributable to the relative abundance of SVs containing different neurotransmitters. 85% to 90% neurons in the mouse brain were glutamatergic while 10%-15% were GABAergic ^171–173^, which can explain our detection of Glu as approximately 10 times that of GABA (Figure 1 B and C). Similarly, cholinergic neurons (5.67−10^5^) ^174^ represented 0.81% of total number of neurons (approximately 70 million) in the mouse brain ^175^, serotonergic neurons (approximately 26,000) for 0.037% of total neurons ^175,176^. Assuming that the content of each neurotransmitter in a single SV is similar, extrapolation from the above data would suggest that approximately 1.3% to 2.15% of neurons in the mouse brain are creatinergic.

To distinguish whether small molecules co-purified with SVs were in the SVs ^143,144^, or that they were just associated with the outside of SVs^154^, we tested the dependence of the presence of these molecules in the SVs on temperature and on the electrochemical gradient of H^+^. Cr was significantly reduced in SVs purified at the room temperature as compared to that immunoisolated at near 0 °C (Figure 1F), supporting the presence of Cr inside, instead of outside, SVs.

Classical neurotransmitters are stored in SVs with an acidic environment inside (pH of 5.6 ∼ 6.4) ^177–179^. To further verify the storage of Cr in SVs and examine the role of H^+^ electrochemical gradient, we applied pharmacological inhibitors during purification ^74,180^. The proton ionophore FCCP (carbonyl cyanide-4-(tri-fluoromethoxy) phenylhydrazone) was used to dissipate H^+^ electrochemical gradient ^180, 181^. FCCP significantly reduced the amount of Cr as well as classical neurotransmitters in SVs (Supplementary Figure 2A-E). The extent of FCCP induced reduction was correlated with the value of pKa or PI (isoelectric point) for different molecule: 5-HT (with pKa predicted to 10 and 9.31, Supplementary Figure 2E) > Cr (PI of ∼7.94, Supplementary Figure 2A) > GABA (PI of 7.33, Supplementary Figure 2C) > Glu (PI of 3.22, Supplementary Figure 2B). Nigericin, a K^+^/H^+^ exchanger which dissipates ΔpH^180,181^, also reduced the amount of Cr and classical neurotransmitters in SVs (Supplementary Figure 2A-E). Furthermore, in the presence of FCCP or nigerin, SV Cr was reduced to a level comparable to that pulled down by control IgG (Supplementary Figure 2A), demonstrating the storage of Cr in SVs was dependent on H^+^ gradient. As a control, the non-neurotransmitter molecule alanine in SVs was not changed by the inhibitors (Supplementary Figure 2F).

### Reduction of SV Cr in mouse mutants lacking slc6a8

SLC6A8, located on the X chromosome, encodes a transporter for Cr and its loss of function (LOF) mutations caused behavioral deficits in humans ^112,113^ and mice ^125–130^. To investigate whether SLC6A8 affects Cr in SVs, we generated slc6a8 knockout (KO) mice. Exon 1 of the SLC6A8 gene was partially replaced with CreERT2-WPRE-polyA by CRISPR/Cas9 (Figure 2A). Examination by reverse polymerase chain reaction (RT-PCR) (Supplementary Figure 3A and 3B) and quantitative real-time reverse PCR (qPCR, Supplementary Figure 3C-D) showed that SLC6A8 mRNA was not detected in either male or female mutants, and significantly reduced in female heterozygous (Slc6a8^+/-^). Consistent with previous reports, the body weights of SLC6A8 KO mice were reduced (Supplementary Figure 4 B and D) ^125,182,183^. Brain weight was not significantly different between SLC6A8 KO mice and WT mice (Supplementary Figure 4A and C).

**Figure 2.**
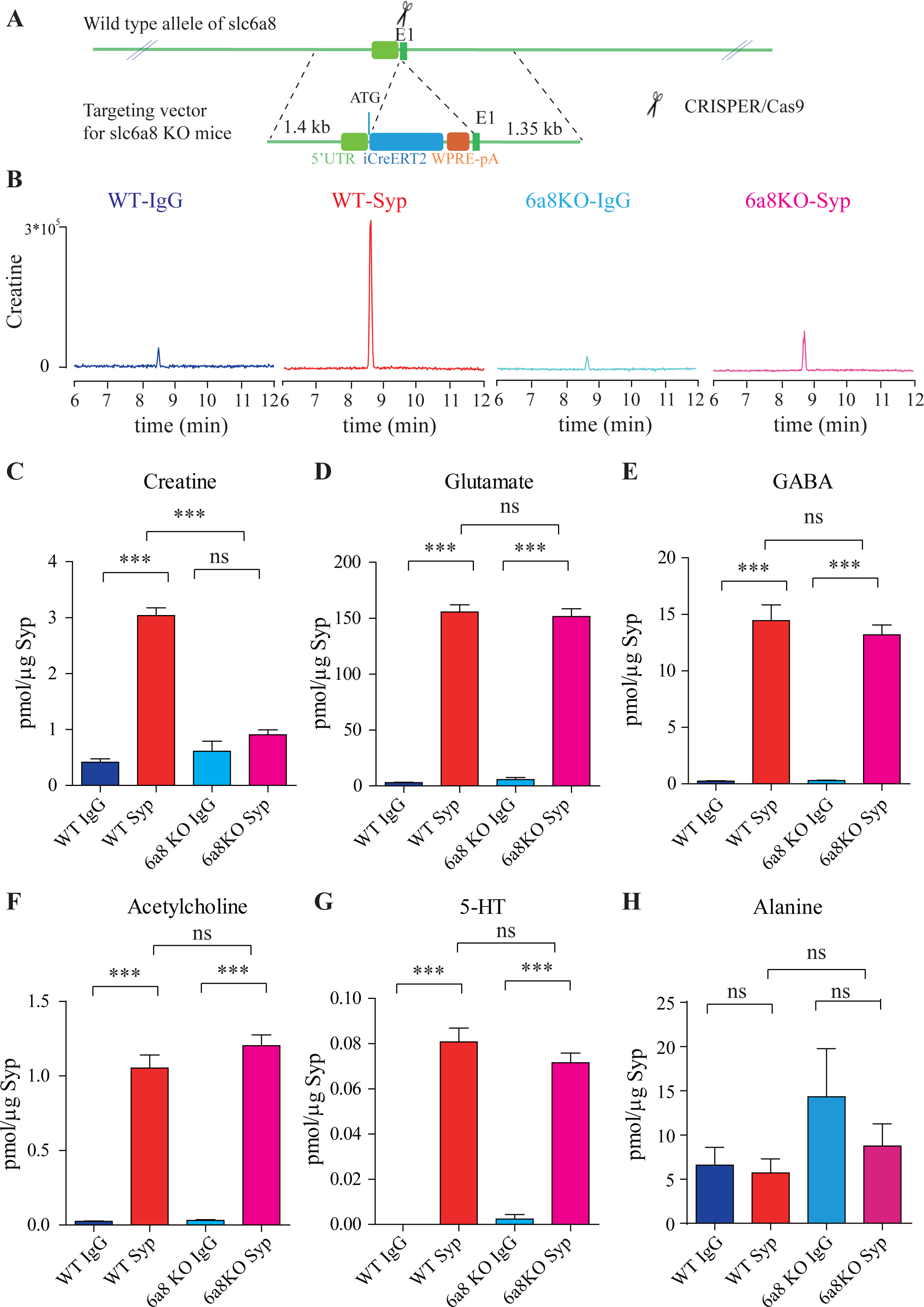
SLC6A8 and Cr in SVs. (A) A schematic illustration of the strategy for generating SLC6A8 knockouts using CRISPR/Cas9. An iCreERT2-WPRE-pA cassette (∼3.5 kb) was inserted immediately downstream of ATG in the SLC6A8 gene, substituting bp 4 to bp 51 in exon 1 (E1). (B) Representative raw traces of Creatine immunoisolated by control IgG from WT mice (blue), the anti-Syp antibody from WT mice (red), IgG from SLC6A8 KO mice (blue) and the anti-Syp antibody from SLC6A8 KO mice (red). (C-H) Quantification of indicated molecules. Note the selective reduction of Cr in SVs from SLC6A8 KO mice. n=14 samples per group. ***, p<0.001, ns, not significant. one-way ANOVA with Tukey’s correction.

When we examined the contents of SVs isolated by the anti-Syp antibody vs the control IgG, significant reduction was only observed for Cr, but not classical neurotransmitters (Figure 2 B-H, Supplementary Figure 5 A-E). While Cr pulled down by IgG was not significantly different between SLC6A8^-/Y^ and SLC6A8^+/Y^ mice, SV Cr purified by the anti-Syp antibody from SLC6A8^-/Y^ was reduced to approximately 1/3 that of the WT (SLC6A8^+/Y^) littermates (n =14, Figure 2 B-C). Compared to the IgG control, Cr in SVs was enriched in WT mice, but not in SLC6A8^-/Y^ mice (Figure 2 B and C). In both SLC6A8^-/Y^ and SLC6A8^+/Y^ mice, classical neurotransmitters in SVs were all enriched as compared to IgG controls (Figure 2 C-G, Supplementary Figure 5 A-D). The amounts of Glu (Figure 2D and Supplementary Figure 5A), GABA (Figure 2E and Supplementary Figure 5B), ACh (Figure 2F and Supplementary Figure 5C) and 5-HT (Figure 2G and Supplementary Figure 5D) in SVs were not different between SLC6A8^-/Y^ and SLC6A8^+/Y^ mice. Molecules not enriched in SVs from WT mice, such as alanine, was also unaffected by SLC6A8 KO (Figure 2H and Supplementary Figure 5E).

It is unlikely that the specific reduction of Cr in SVs from SLC6A8 KO mice was due to technical artifacts. First, the possibility of less SVs obtained from SLC6A8 KO mice were precluded by immunoblot analysis, as assessed by SV markers Syp, Syt and H^+^-ATPase (Supplementary Figure 6A). Second, data collected by high resolution MS (Q Exactive HF-X, Thermoscientific) also revealed selective decrease of SV Cr (m/z=132.0576) from SLC6A8 KO mice (n=8, Supplementary Figure 6 B-G), as quantified by the peak area. Peak areas for Glu (n=8, Supplementary Figure 6B, m/z=148.0604), GABA (n=8, Supplementary Figure 6C, m/z=104.0712), ACh (n=8, Supplementary Figure 6D, m/z = 146.1178) and alanine (n=8, Ala, Supplementary Figure 6E, m/z=90.055) were not significantly different between SVs immmunoisolated with the anti-Syp antibody and control IgG from WT and Slc6a8 KO mice. However, peak areas (Supplementary Figure 5G) and amplitude of Cr (n=8, Supplementary Figure 5F) signal were significantly increased in SVs from WT mice (anti-Syp antibody vs IgG), but not that from SLC6A8 KO mice.

### Reduction of SV Cr in mouse mutants lacking AGAT

AGAT is the enzyme catalyzing the first step in Cr synthesis ^82,184^ and its absence also led to Cr deficiency in the human brain and mental retardation ^132,133^. To investigate requirement of AGAT for SV Cr, we utilized AGAT ‘knockout-first’ mice (Figure 6A) ^185^. The targeting cassette containing Frt (Flip recombination sites)-flanked EnS2A, an IRES::lacZ trapping cassette and a floxed *neo* cassette were inserted downstream of exon 2 to interfere with normal splicing of AGAT pre-mRNA. Examination by RT-PCR (Supplementary Figure 7A) and quantitative RT-PCR (Supplementary Figure 7B) showed reduction of AGAT mRNA in AGAT^+/-^ mice and absence of AGAT mRNA in AGAT^-/-^ mice. Body weight (Supplementary Figure 8B), but not brain weight (Supplementary Figure 8A), of AGAT^-/-^ mice were lower than both AGAT^+/+^ and AGAT^+/-^ mice, which were similar to SLC6A8 KO mice.

Immunoblot analysis showed SVs purified from the brains were not significantly different among AGAT^+/+^, AGAT^+/-^ and AGAT^-/-^ mice (Supplementary Figure 9 A-D), as supported by quantitative analysis of Syp, Syt and H^+^-ATPase (n=20, with 2 repeats for 10 samples).

We analyzed small molecules present in SVs from AGAT^+/+^, AGAT^+/-^ and AGAT^-/-^ mice. Cr was significantly enriched in SVs from all three genotypes as compared to the IgG control (Figure 3A and 3B). However, the level of Cr from AGAT^-/-^ mice was significantly lower than those from AGAT^+/+^ and AGAT^+/-^ mice (n=10, Figure 3A and B).

**Figure 3.**
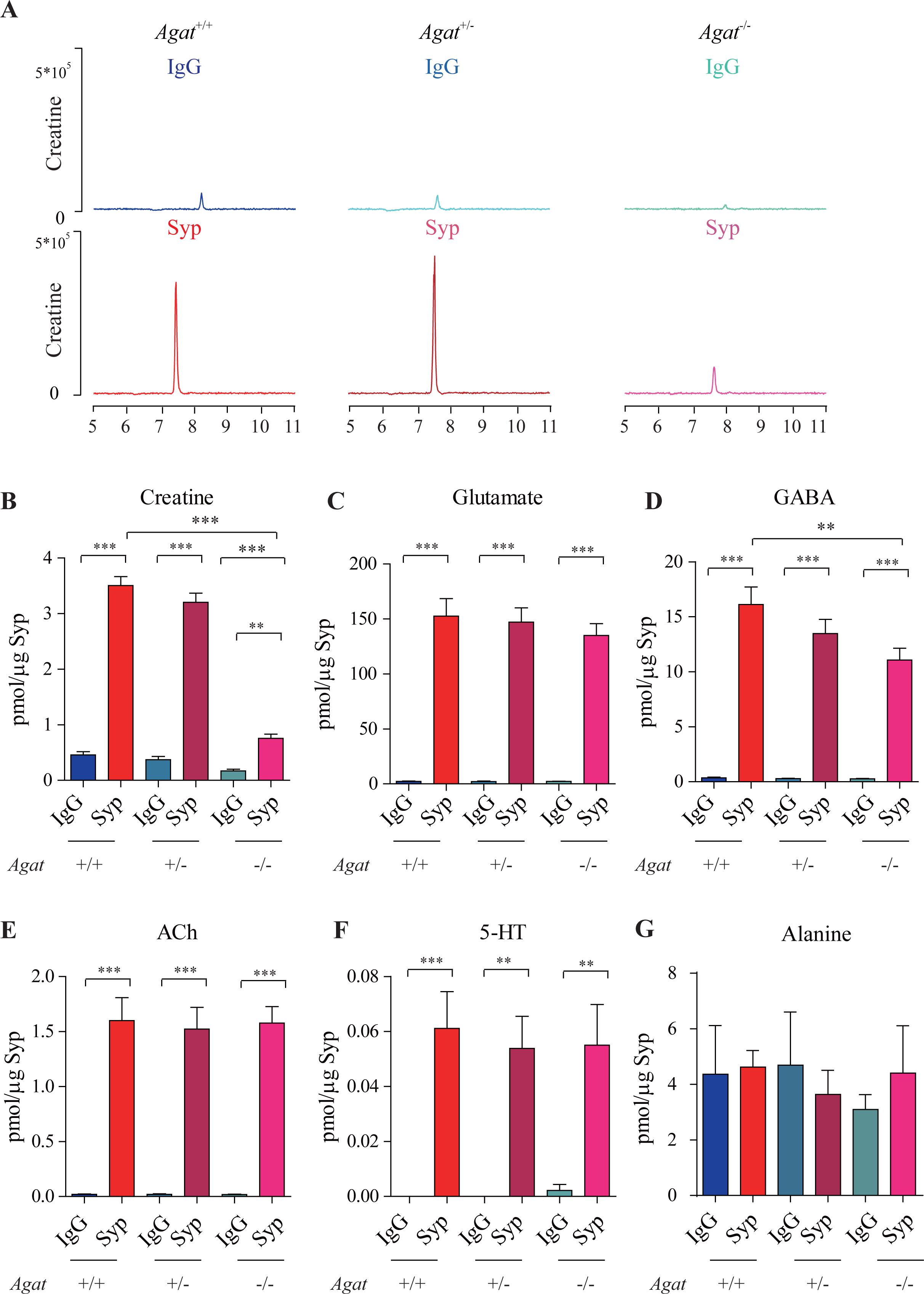
Reduction of SV Cr in AGAT knockout mice. (A) Representative raw traces of Creatine, pulled down by IgG or anti-Syp from AGAT^+/+^, AGAT^+/-^ and AGAT^-/-^ mice. (B-G) Quantification of indicated molecules. Cr was significantly decreased in AGAT^-/-^ mice as compared to Cr in AGAT^+/+^ or AGAT^+/-^ mice (B). GABA was significantly decreased in SVs from AGAT^-/-^ mice as compared to AGAT^+/+^ mice (D) but the difference was smaller than that of Cr. Glu (C), ACh (E), 5-HT (F) and alanine was not different among AGAT^+/+^, AGAT^+/-^ and AGAT^-/-^ mice. n=10 samples per group. ***, p<0.001, ns, not significant. one-way ANOVA with Tukey’s correction.

Glu (Figure 3C), ACh (Figure 3E) and 5-HT (Figure 3F) were all enriched in SVs (as compared to IgG controls), and not significantly different among AGAT^+/+^, AGAT^+/-^ and AGAT^-/-^ mice. GABA in SVs from AGAT^-/-^ mice was also decreased from AGAT^+/+^ mice by 30%, to an extent less than that of Cr (78.4%). Alanine was not different among three genotypes of mice (n=6, Figure 3G). Thus, Cr and GABA, but not other neurotransmitters, in SVs were reduced in AGAT KO mice.

### Pattern of SLC6A8 expression indicated by knockin mice

We generated SLC6A8-HA knockin mice by CRISPR/Cas9. Three repeats of the hemagglutinin (HA) tag ^186^, the T2A sequence ^187–189^ and CreERT2 ^190–193^ were inserted in-frame at the C terminus of the SLC6A8 protein (Figure 4A).

**Figure 4.**
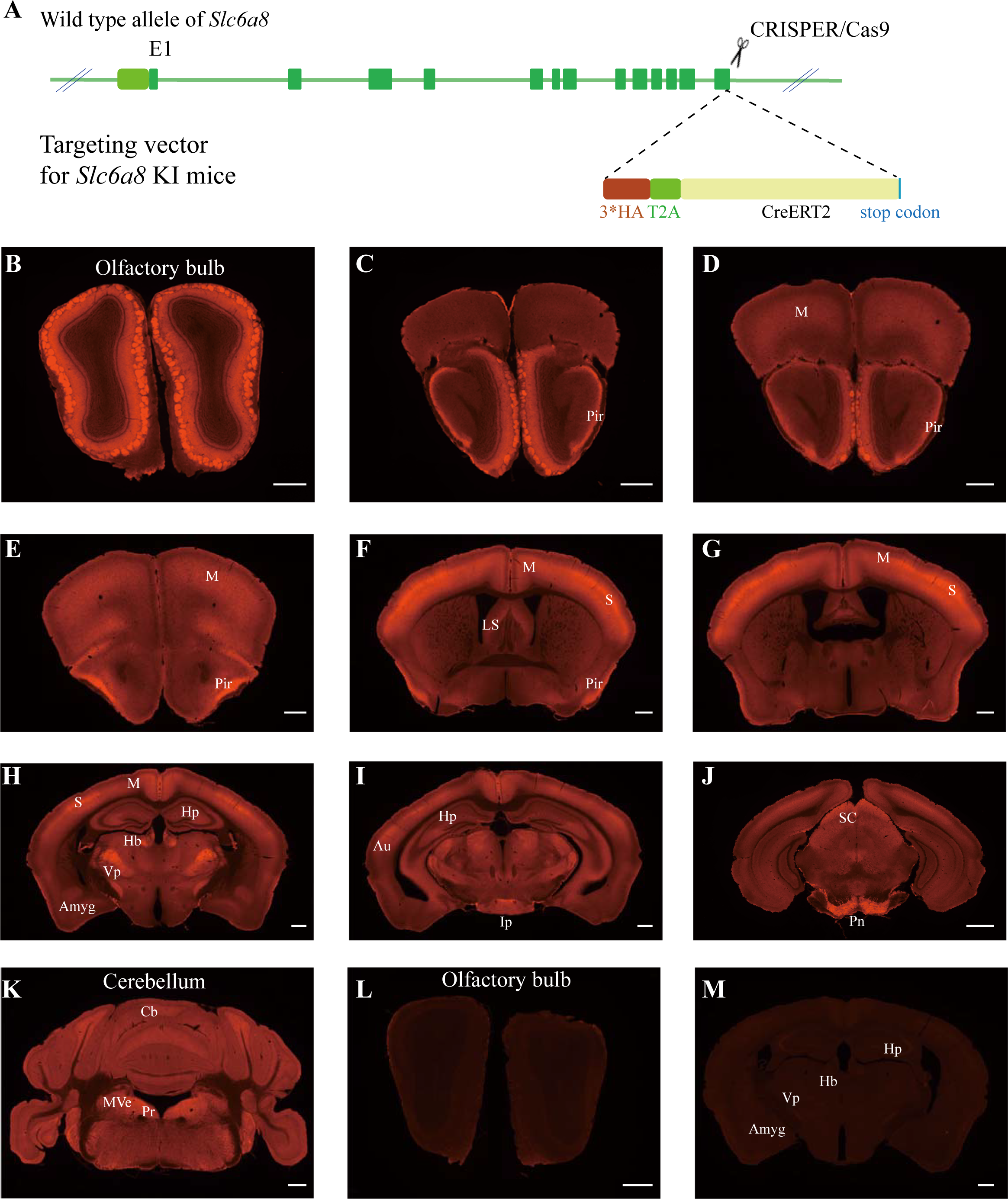
Expression pattern of SLC6a8. (A) A diagram illustrating the generation of SLC6A8-HA KI mice. 3*HA, T2A and CreERT2 were inserted in-frame, before the stop codon, to the C terminus of SLC6A8 protein. (B-K) Low magnification photomicrographs of coronal sections immunohistochemically labelled with the anti-HA antibody in SLC6A8-HA mice. (I and M) immunostaining with the anti-HA antibody in control WT mice. Abbreviations: Pir, piriform cortex; M, motor cortex; LS: lateral septum; Hp: hippocampus; Hb, habenular nucleus; Vp: ventral posterior nucleus of thalamus; Au auditory cortex; Amyg: amygdala; Ip, interpeduncular nucleus; Pn; pontine nucleus; Cb, cerebellum; Pr, prepositus; SC, Superior colliculus; MVe, medial vestibular nucleus. Scale bar: 500 μm.

To examine the expression pattern of SLC6A8, we performed immunocytochemistry with an antibody against the HA epitope in SLC6A8-HA and WT mice. SLC6A8-HA mice showed positive signals in the olfactory bulb (Figure 4B), the piriform cortex (Figure 4 C-F), the somatosensory cortex (Figure 4 F and G), the ventral posterior thalamus (Figure 4H), the interpeduncular nucleus (Figure 4I) and the pontine nuclei (Figure 4J). In addition, moderate levels of immunoreactivity were observed in the motor cortex (Figure 4 D-H), the medial habenular nucleus (Figure 4H), the hippocampus (Figure 4H) and the cerebellum (Figure 4K). These results were consistent with previous reports ^109,111^. WT mice were negative for anti-HA antibody staining (Figure 4 L and M).

### Ca2+ dependent release of Cr upon stimulation

Classical neurotransmitters are released from the SVs into the synaptic cleft in a Ca^2+^ dependent manner after stimulation. For example, high extracellular potassium (K^+^) stimulated Ca^2+^ dependent release of Glu, GABA and other neurotransmitters in brain slices ^194–198^.

300 μm thick coronal slices of the mouse brain within 1∼2 millimeters (mm) posterior to the bregma were used, because the cortex, the thalamus, the habenular nucleus and the hippocampus were positive for SLC6A8 (cf., Figure 4H). We monitored the effect of K^+^ stimulation by recording neurons in the slices. Immediately after K^+^ stimulation, pyramidal neurons in the CA1 region of the hippocampus were depolarized, firing a train of action potentials and reaching a large depolarization plateau in less than 1 minute (min) (Figure 5A). K^+^-induced depolarization persisted for several mins before returning to the baseline and being washed within 10 mins. Thus, superfusates in 1-min fraction at the time points of 1.5 min before (control) and after K^+^ stimulation, and 10 mins after the wash were collected (Figure 5A), and the metabolites in the superfusates were analyzed by CE-MS.

**Figure 5.**
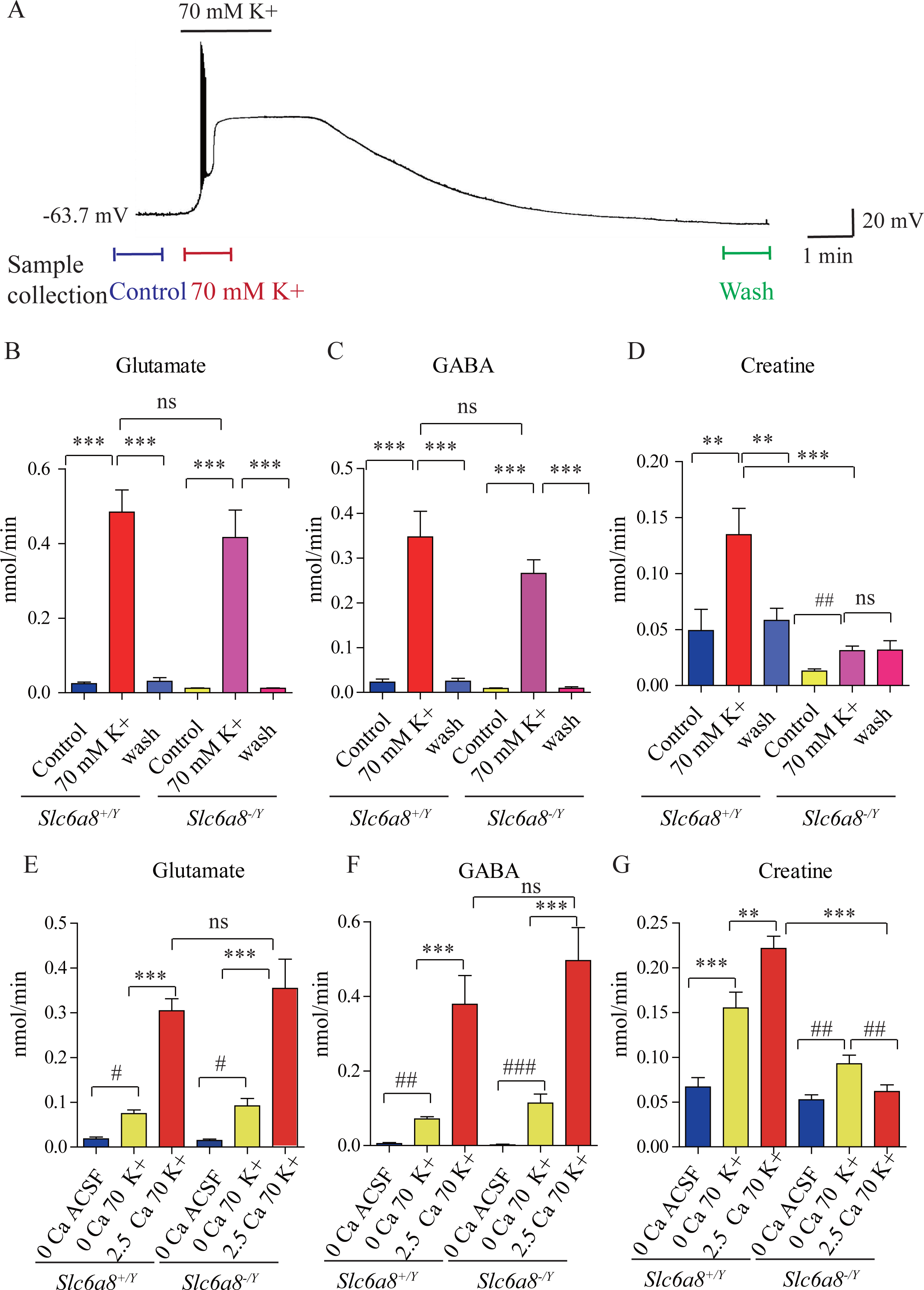
Cr release in brain slices from WT and SLC6a8 knockout mice. (A) Neuronal depolarization induced by 70 mM K^+^ and time points for collecting the release sample. “Control” samples were collected 1.5 min to 0.5 min before K^+^ stimulation, “70 mM K^+^” ACSF samples were collected during 70 mM K^+^ stimulation, and “wash” samples were collected 10 mins after washout with ACSF. Efflux of Glu (B), GABA (C) or Cr (D) from WT or SLC6A8 KO male mice (n=7 samples per group). Note that a small amount of Cr released in SLC6A8 KO mice did not return to the baseline after 10 mins of washing. (E-G) Ca^2+^ dependent release of Glu, GABA and Cr in WT and SLC6A8 KO mice (n=5 samples per group). ***, p<0.001, ns, not significant. one-way ANOVA with Tukey’s correction. #, p<0.05, ##, p<0.01, paired t test.

**Figure 6.**
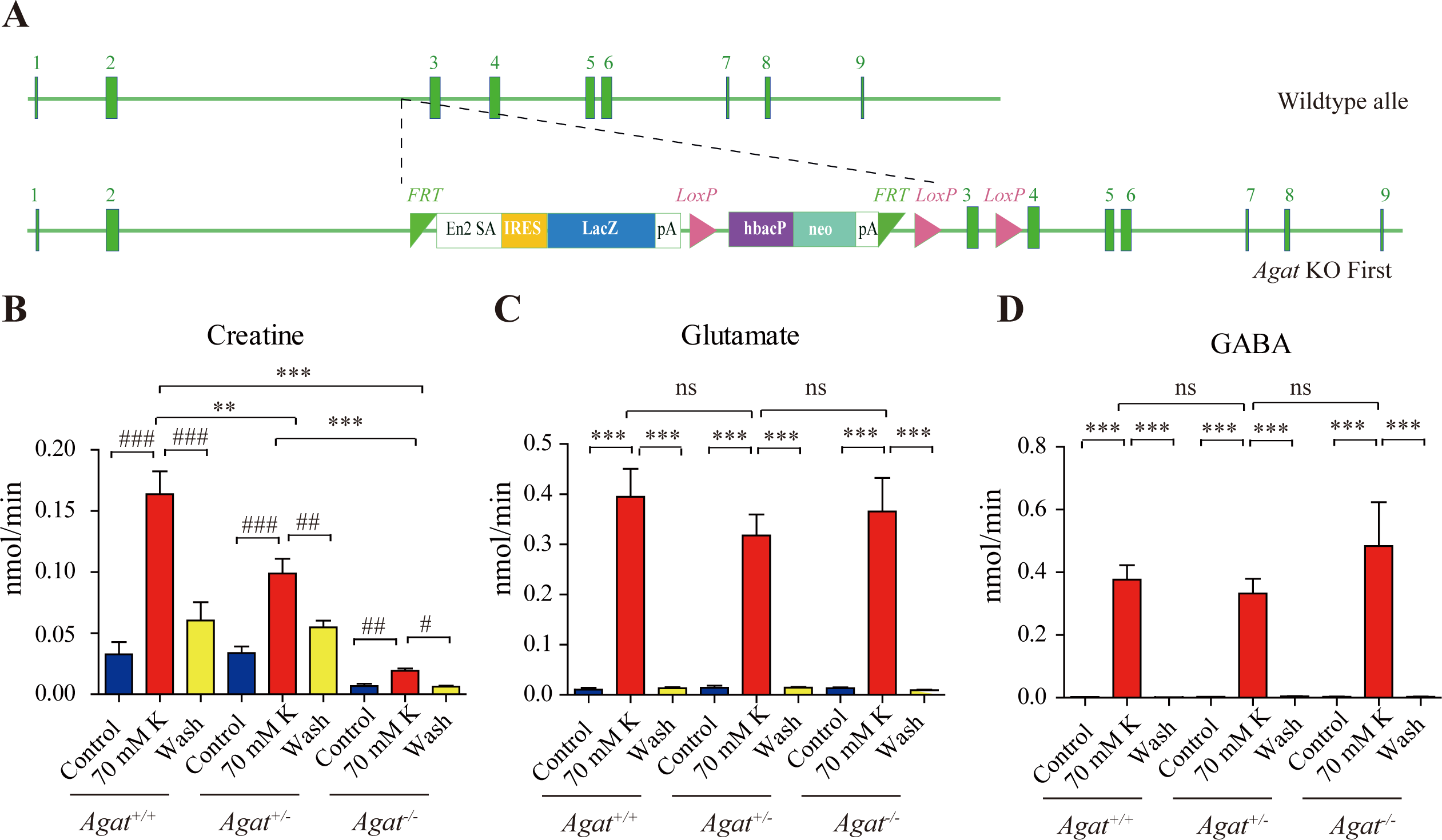
Cr release in WT and AGAT knockout mice. (A) A schematic diagram illustrating the strategy of AGAT knockout first. With the AGAT gene (also known as GATM) shown in the upper part, and the gene targeting strategy in the lower part. The homologous arm is approximately 10 kb. A targeting cassette, containing Frt-flanked lacZ and neomycin, was inserted downstream of exon 2. At the same time, exon 3 of AGAT was flanked by two loxP sites. K^+^ induced release of Glu (C) and GABA (D) were not significantly different among AGAT^+/+^, AGAT^+/-^ and AGAT^-/-^ mice, whereas that of Cr (B) was significantly lower in AGAT^-/-^ mice than those in AGAT^+/+^ and AGAT^+/-^ mice.

In the presence of Ca^2+^, depolarization with elevated extracellular K^+^ led to robust release of Glu and GABA in slices from WT (SLC6A8^+^/Y) mice (n=7 per group, Figure 5 B and C). After 10 mins wash, levels of Glu and GABA returned to the baseline (Figure 5 B and C). In the presence of Ca^2+^, depolarization with elevated K^+^ led to robust release of Cr. Extracellular Cr returned to the baseline after 10 min wash (Figure 5D). For quantification, the stimulated releases of metabolites were calculated by subtracting the basal levels from the total releases in response to K^+^ stimulation. In the presence of Ca^2+^, K^+^ stimulation induced the efflux of Glu, GABA and Cr at 0.46, 0.33 and 0.086 nmol/min, respectively (n=7 per group) (Figure 5 B, C, D). From the detection limits of ACh and 5-HT in our system, we inferred that the efflux rate for ACh was lower than 0.001 nmol/min and that for 5-HT lower than 0.003 nmol/min. The efflux rate for Cr in brain slices is lower than those of Glu and GABA, but higher than those for ACh and 5-HT.

Ca^2+^ dependence of transmitter release was examined by comparing responses to ACSF without Ca^2+^ or elevated K^+^ (supplemented with 1 mM EGTA), elevated extracellular K^+^ in the absence of Ca^2+^ (supplemented with 1 mM EGTA), or K^+^ in presence of 2.5 mM Ca^2+^ (Figure 5 E-G, n=5 per group). In the absence of Ca^2+^, elevated K^+^ stimulated the release of a small but significant amount of Glu and GABA, with efflux rates at 0.056 nmol/min and 0.066 respectively (Figure 5 E, F). In the presence of 2.5 mM Ca^2+^, elevated K^+^ further augmented the release of Glu and GABA by 5 to 6 times, confirming previously reported Ca^2+^ dependent release of neurotransmitters in response to depolarization ^194,196^.

Cr was also released both in a Ca^2+^ dependent and a Ca^2+^ independent manner (Figure 5G). More Cr was released in response to K^+^ stimulation in the presence of 2.5 mM Ca^2+^ than that in the absence of Ca^2+^. These results demonstrate Ca^2+^ dependent release of Cr upon stimulation.

### Reduced Cr release in SLC6A8 and AGAT mutant mice

We examined whether SLC6A8 KO affected K^+^-induced release of Cr. While Glu and GABA were released in slices from SLC6A8 KO (SLC6A8^-^/Y) mice at levels not significantly different from those of WT mice (Figure 5 B, C), release of Cr in response to K^+^ stimulation was significantly reduced in SLC6A8^-/Y^ mice as compared to SLC6A8^+/Y^ mice (Figure 5D). The basal level of Cr in SLC6A8 KO mice was lower than that of WT mice. In addition, K^+^ stimulation induced release of Cr persisted to some extent even after 10 mins of washout (Figure 5D), possibly due to the inability of presynaptic terminals in SLC6A8 KO mice to reuptake Cr in the synaptic cleft (Figure 8).

Experiments with slices from brains of SLC6A8 KO (SLC6A8^-^/Y) mice showed that Ca^2+^ dependent release of either Glu or GABA was not affected by the genotype of SLC6A8 (Figure 5 E and F). By contrast, Ca^2+^ dependent release of Cr was abolished in SLC6A8^-^/Y slices. Interestingly, Ca^2+^ independent release of Cr was reduced by a third, but did not reach statistical significance, in SLC6A8^-^/Y slices. In the absence of Ca^2+^, the basal level of Cr was not changed in SLC6A8 KO mice. Taken together, these results indicate that there is Ca^2+^ dependent release of Cr upon stimulation and that SLC6A8 is required specifically for Ca^2+^ dependent release of Cr, but not for Ca^2+^ dependent release of other neurotransmitters such as Glu and GABA, or for Ca^2+^ independent release of Cr. Knockout of AGAT (Figure 6A) selectively reduced K^+^ evoked release of Cr, but not those of Glu or GABA (n=5 per group, Figure 6 B, C, D). Although K^+^ stimulation still elicited Cr release from brain slices of AGAT^+/-^ mice, the efflux rate in AGAT^-/-^ mice was reduced to less than 10% that in AGAT^+/+^ mice and 20% that in AGAT^+/-^ mice.

### Cr inhibition of neocortical neurons

Our own data (Figure 4H) and previous reports ^109,111^ have shown SLC6A8 in the neocortex, with dense SLA6A8-HA immunoreactive fibers in layer 4 (Supplementary Figure 10A) ^199,200^. Layer 5 neurons in the somatosensory cortex have been reported to express SLC6A8 previously ^109,111^. To investigate electrophysiological effects of Cr, we performed whole cell patch clamp recordings from the pyramidal neurons in layer 4/5 of the somatosensory cortex (Figure 7 A and B, Supplementary Figure 10).

**Figure 7.**
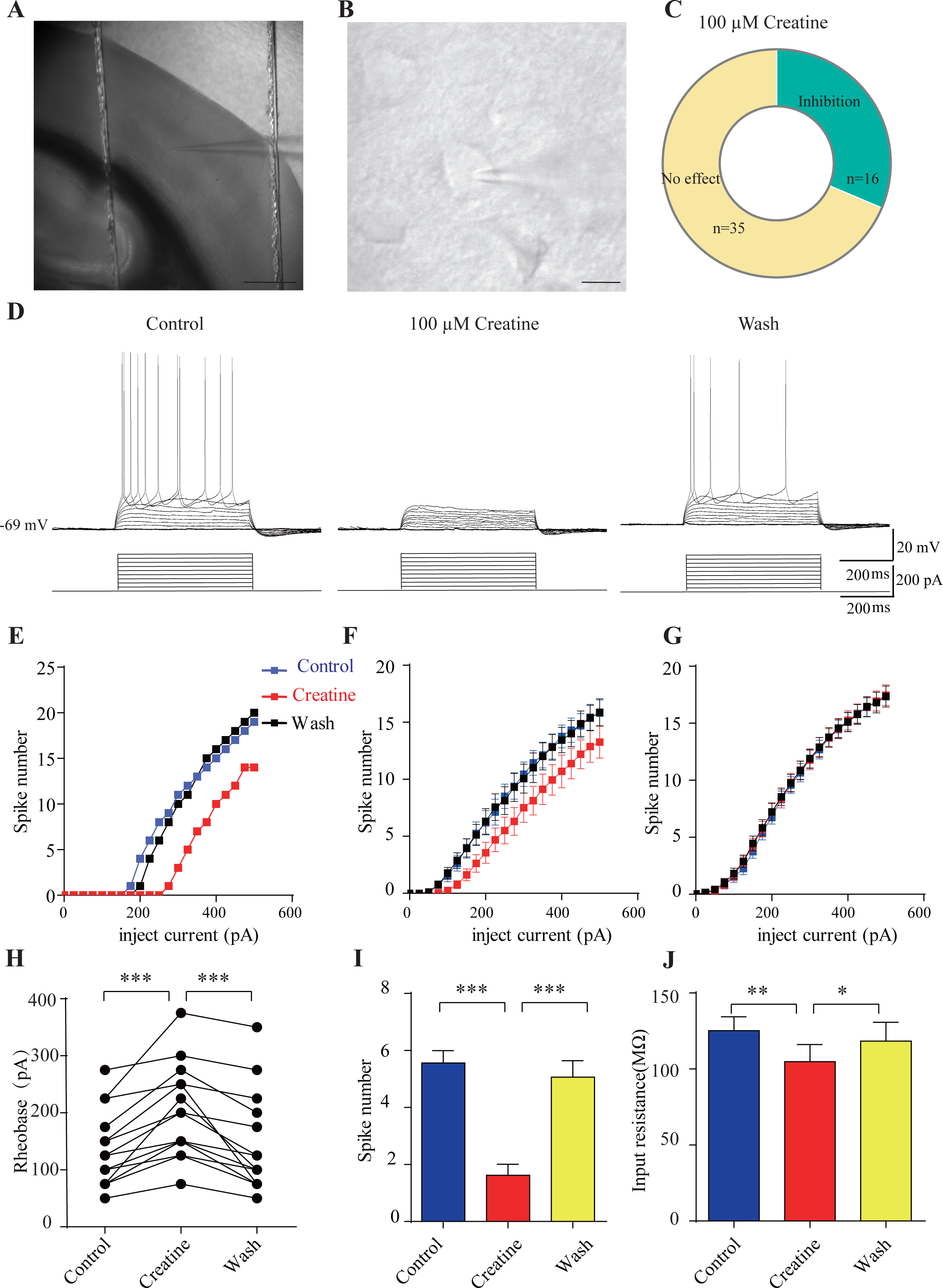
Inhibitory effects of Cr on cortical neurons. (A) A photograph showing recording at layer 4 in the somatosensory cortex. Scale bar: 10 μm. (B) Patch-clamp recording of a pyramidal neuron. Scale bar: 10 μm. (C) Ratios of Cr-responsive and -unresponsive neurons in the region. (D) Representative raw electrophysiological traces showing inhibition of evoked firing by Cr, with the lower panel showing the stimulus protocol. (E) Evoked spike numbers in response to different current injections from (D). (F) Relationship between evoked spike numbers and different current injections to neurons that were inhibited by Cr (n=16). (G) The same for Cr-unresponsive neurons (n=35). (H) Rheobase for Cr responsive neurons. (I) Evoked spike number when these neurons were injected with current of rheobase + 50 pA. (J) input resistance. *, p<0.05; **, p<0.01; ***, p<0.001, paired t test.

**Figure 8.**
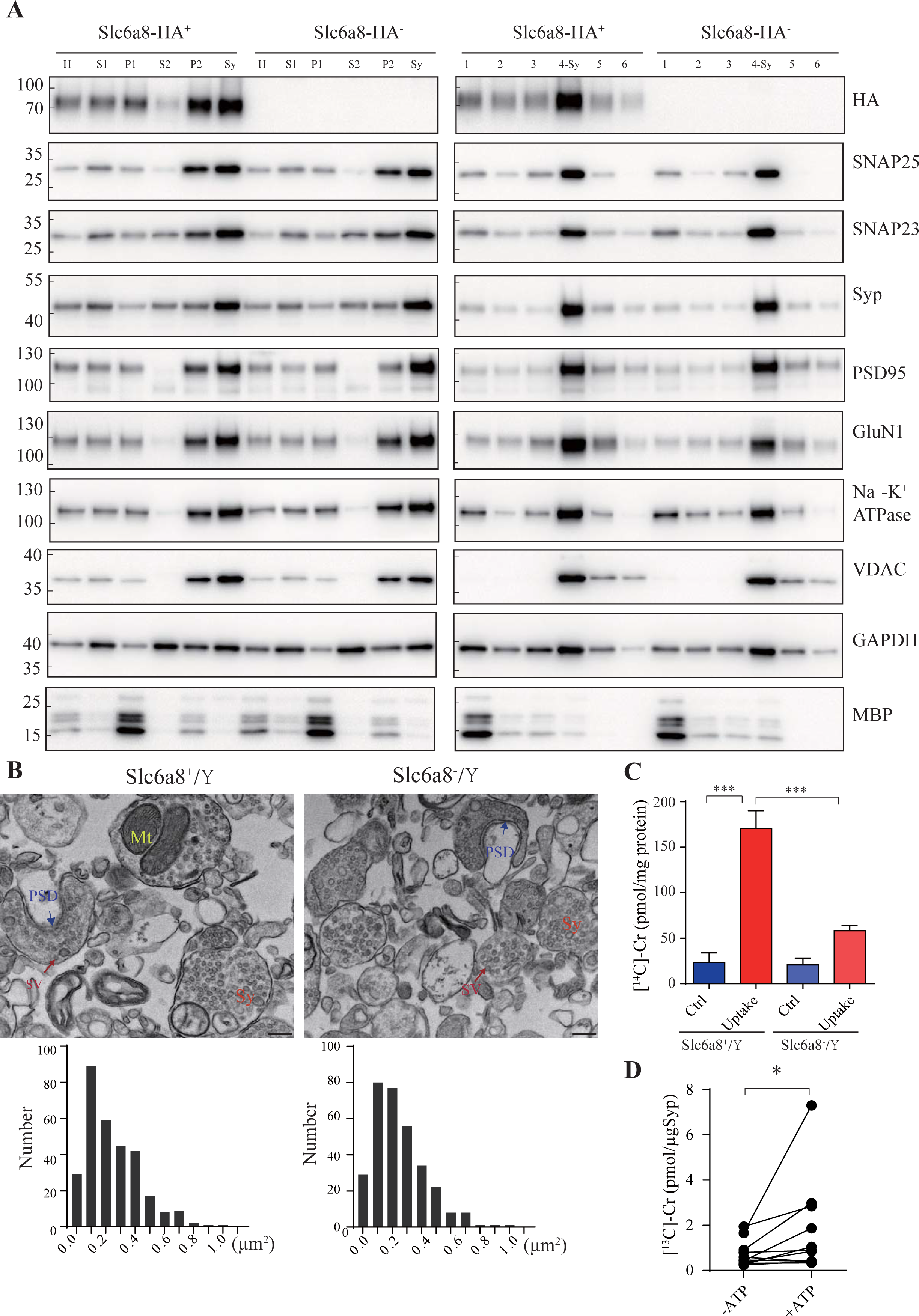
Cr uptake into synaptosomes and SVs. (A) Markers for subcellular organelles detected in synaptosomes prepared from WT mice or mice with Slc6a8 gene fused in frame to the HA epitope. SLC6A8-HA was enriched in synaptosomes (Sy or 4-Sy) or crude synaptosomes (P2) (see methods), as were other markers for the subcellular organelles in synaptosomes but not the marker for myelin (MBP). (B) Representative electron micrographs and histograms of size distribution in synaptosomes isolated from WT (Slc6a8^+^/Y) and *Slc6a8* KO (Slc6a8^-^/Y) mice by Ficoll density-gradient centrifugation. Sy, synaptosome; SV, synaptic vesicles; Mt, mitochondria; PSD, post-synaptic density. Bar, 20 nm. (C) Cr uptake into synaptosomes (n=5 per group). The two left columns were results from WT mice and the two right columns from Slc6a8 knockout mice. The control baseline was [^14^C]-Cr uptake at 0 °C at 10 mins (*205*). Cr uptake into synaptosomes at 37 °C measured at 10 mins was observed in WT synaptosomes. Uptake into *Slc6a8* knockout synaptosomes was significantly reduced when compared to the WT synaptosomes. ***, p<0.001, one-way ANOVA with Tukey’s correction. (D) uptake of [^13^C]-Cr into immunoisolated SVs in the presence or absence of ATP. (n=11 samples per group)*, p<0.05, paired-t test.

Medium-sized pyramidal neurons (Figure 7B) with a membrane capacitance (Cm) of 114.96 ± 3.92 pF (n=51, Supplementary Figure 11A) were recorded. These neurons exhibited regular firing patterns ^201,202^ in response to depolarization current injection (Figure 7D and Supplementary Figure 11A) with moderate maximal evoked spiking frequencies of 10-30 spikes per 500 milliseconds (ms) (Figure 7 E-G), increasing of inter-spike intervals during depolarizing steps (Figure 7D and Supplementary Figure 11A), high action potential amplitude (81.64 ± 1.06 mV, Supplementary Figure 12E) and large spike half-width (1.12 ± 0.031 ms, Supplementary Figure 12F).

Cr was bath applied only after the evoked firing pattern reached a steady state. Of the 51 neurons, 16 were inhibited by 100 μM creatine (Figure 7 B-F). Fewer spikes were evoked in Cr responsive neurons in response to depolarizing current injections during Cr application (25 pA step, 500 ms) (Figure 7 D-F). The inhibitory effect of Cr was reversible (Figure 7 D-F), typically observed within 2-3 min following Cr application (with maximal effect from 2 to 8 mins) and disappeared after 10 to 25 min washout. This could be repeated by a second application of Cr. The rheobase, defined as the minimal electrical current necessary to elicit an action potential, was increased during bath application of Cr (Figure 7 H). The inhibitory effect was most obvious at near spike threshold. When a neuron was depolarized with a current of 50 pA above rheobase, the number of evoked spikes was decreased dramatically during Cr application (Figure 7I). Cr also mildly inhibited the input resistance (Figure 7J), slightly hyperpolarized resting membrane potential (Supplementary Figure 12C) or reduced amplitude of afterhyperploarization (AHP) followed by the first evoked action potential (Supplementary Figure 12G). The spike threshold (Supplementary Figure 12D), amplitude (Supplementary Figure 12E) and half width (Supplementary Figure 12F) were not changed by Cr.

The remaining 35 neurons were not responsive to Cr (Figure 7C and G, Supplementary Figure 11). Cr did not change electrical parameters tested, including evoked firing rates (Figure 7G, Supplementary Figure 11 A, B and D), rheobase (Supplementary Figure 11C), resting membrane potential (Supplementary Figure 12C), spike threshold amplitude (Supplementary Figure 12E) and half width (Supplementary Figure 12F). In addition, electrical properties of responsive neurons and unresponsive neurons were not significantly different. With the limited number of neurons recorded, the ratio of responsive neurons appeared higher in layer 4 or border of layer4/5, than the deeper layer in layer 5 (Supplementary Figure 10).

### SLC6A8 dependent uptake of Cr into the synaptosomes

Along with enzymatic degradation, re-uptake by transporters serves as an important way to remove neurotransmitters released into the synaptic cleft. As synaptosomes contain the apparatus for neurotransmission, they are often used for studying uptake of neurotransmitters ^203^.

To investigate whether Cr uptake into synaptosomes required SLC6A8, we first examined whether SLC6A8 was present in synaptosomes. Using Slc6a8-HA knockin mice and an anti-HA antibody, we found that SLC6A8-HA was present and enriched in crude synaptosomal fraction (P2 fraction in Figure 8A, enrichment score: P2/H = 1.76 ± 0.15, n = 4) and synaptosomal fraction prepared using a discontinuous Ficoll gradient (Sy and 4-Sy fractions in Figure 8A, enrichment score: Sy/H = 2.02 ±0.14, n = 4). The integrity of synaptosomes was confirmed by multiple markers of the synaptosomes ^203^, including the presynaptic membrane marker SNAP25 ^204^ and the SV marker Syp (synaptophysin Syp) ^146–149^ in presynaptic terminals, the postsynaptic density marker PSD95 ^165,166^ and the postsynaptic membrane protein GluN1 ^205^, the synaptic membrane protein SNAP23 ^162–164^, the plasma membrane marker Na^+^-K^+^-ATPase ^206–208^, and the mitochondria marker VDAC ^203^. These were all enriched in our synaptosomal preparations. The cytosol marker GAPDH was also present in synaptosomes, whereas the oligodendrocyte marker MBP was nearly absent, suggesting myelin pollution was largely avoided ^203^.

We have also used EM to confirm the quality of our synaptosome preparations. As reported previously ^203,209^, synaptosomes were composed of membrane bounded structures (Sy in Figure 8B) filled with synaptic vesicles (SV in Figure 8B), sometimes with a segment of postsynaptic membrane along with the postsynaptic density (PSD in Figure 8B) and mitochondria (Mt in Figure 8B). The sizes of synaptosomes from WT mice and *Slc6a8* knockout mice were similar, with areas of 0.245 ± 0.01 μm^2^ (n=302 particles) and 0.247 ± 0.01 μm^2^ (n=317 particles), respectively (Figure 8B).

We then examined whether SLC6a8 participated in Cr uptake into the synaptosomes. A mixture of 18 μM [^14^C]-Cr (with a total radioactivity of 0.4 μCi) and 5 μM Cr was used, and uptake at 0 °C measured at 10 mins was the baseline ^210^. Cr uptake into synaptosomes from WT mice was stimulated approximately sevenfold at 37 °C (Uptake, Figure 8C) as compared with 0 °C (Ctrl, Figure 8C). Cr uptake into synaptosomes from SLC6a8 knockout mice was less than 3 times as compared to its control, and was decreased to approximately 1/3 of that of WT mice (Figure 8C). Thus, SLC6A8 is necessary for uptake of Cr into the synaptosomes.

### Cr uptake into SVs

Classical neurotransmitters were taken up in SVs in an ATP-dependent manner^144,159,181,210,211^. We examined whether Cr could be transported into SVs.

We used 10 μg anti-Syp antibody to purify SVs from mouse brains. Purified SVs were preincubated for 30 min to allow sufficient leakage of endogenous Cr, before being mixed with 1 mM [^13^C]-Cr in the presence or absence of 4 mM ATP and placed at 25 L for 10 mins to allow adequate uptake. The SV content of [^13^C]-Cr was then examined by CE-MS and high-performance liquid chromatography-mass spectrometry (HPLC-MS). Significantly more [^13^C]-Cr were taken up by SVs in the presence of ATP, with about 10.3 pmol [^13^C]-Cr transported into SVs (1.03 pmol/μg α-Syp or transportation rate of 0.103 pmol/min, n=11, Figure 8D).

In summary, Cr could be transported into SVs in ATP-dependent manner. At this point, we do not know what is the transporter(s) on the SVs for Cr uptake. SLC6A8 is only found in plasma membrane, not on SVs, and is not a candidate for Cr uptake into SVs.

## Discussion

While no neurotransmitter has been proven in a single paper, supportive evidence suggesting Cr as a possible new neurotransmitter has been presented here to the extent of any single previous papers.

At various times and by different researchers, taurine ^46,212^, proline ^73^, D-aspartic acid ^213^, hydrogen sulfide ^214^, agmatine ^215^, DOPA ^216^, estradiol ^217^, β-alanine ^218^, protons ^219^ have been suspected as neurotransmitters, but they do not meet all the criteria. Some of the suspected molecules can be released upon stimulation or removed by transporters. Often, they have not been reproducibly found in SVs ^74^.

Our discovery of Cr in SVs significantly raised the priority of testing the candidacy of Cr, and our further investigations have led to more evidence suggesting Cr as a neurotransmitter: 1) Cr is stored in SVs; 2) Ca^++^ dependent release of Cr upon stimulation has been observed; 3) both Cr storage in SVs and Cr release are reduced when either the gene for SLC6A8 or the gene for AGAT was deficient; and 4) Cr inhibits activities of pyramidal neurons in the neocortex; 5) Cr uptake into synaptosomes requires SLC6A8; 6) Cr uptake into SVs was ATP-dependent.

Of the above results, 1, 3, 4 and 6 are reported for the first time in this paper. Furthermore, we have demonstrated that detection of Cr in SVs was lower than those for Glu and GABA, but higher than those for ACh and 5-HT, placing Cr at a level in the middle of known central transmitters (Figures 1-3). The storage of Cr in SVs is dependent on preserved H^+^ gradient (Supplementary Figure 2) and Cr can be transported into SVs (Figure 8D).

There was a single previous report of Ca^++^ dependent release of [^3^H]Cr and endogenous Cr in response to electrical stimulation ^220^. We now provide evidence that Cr was released in response to extracellular K^+^ stimulation (within 1-2 min) (Figure 5-6). Furthermore, Cr release was reduced when either the Slc6A8 or AGAT gene was removed (Figures 5 and 6). Although the Ca^2+^ dependent component of K^+^ evoked Cr release was smaller as compared to those of Glu and GABA, it nevertheless existed and was totally abolished by Slc6A8 knockout (Figure 5). The reported electrically evoked Cr release showed more Ca^2+^ dependence ^220^. Taken together, our data and previous report ^220^ supported a role of Cr as a neurotransmitter. Our observation of extremely low efflux rates of 5-HT or ACh may have arisen from very limited numbers of cholinergic^174^ or serotoninergic neurons^176^ in the sliced sections and rapid enzymatic degradation of these neurotransmitters.

Cr uptake from the extracellular space into the cells was reported twice previously, once with brain slices showing sodium dependent uptake of [^3^H]Cr ^220^ and once with synaptosomes ^221^. Our new results have not only replicated the synaptosome Cr uptake experiment but also shown the requirement of SLC6A8, a membrane transporter expressed in synaptosomes (Figure 8A), for Cr uptake into synaptosomes. Transportation of Cr into synaptosomes by Slc6A8 may function for both the clearance of Cr from the synaptic cleft and recycling of Cr into SVs residing in neurons (Figure 8B, Figure 6D).

In summary, in addition to confirming and extending previous results which have stood alone for more than a decade without replication or follow-up, we have obtained entirely new results suggesting the candidacy of Cr as a neurotransmitter. We discuss below the criteria for a neurotransmitter, Cr as a neurotransmitter, and the implications of Cr as a neurotransmitter.

### Criteria of a neurotransmitter

The criteria for establishing a non-peptide small molecule as a neurotransmitter have varied from time to time and from author to author.

Some textbooks simply state that a neurotransmitter is stored presynaptically, released upon stimulation and active on postsynaptic neurons. The details of these 3 criteria can vary. For example, one textbook stipulates that “the substance must be present within the presynaptic neuron; the substance must be released in response to presynaptic depolarization, and the release must be Ca^2+^ dependent; specific receptors for the substance be present on the postsynaptic cell” ^67,68^. Another states that “the molecule must be synthesized and stored in the presynaptic neuron; the molecule must be released by the presynaptic axon terminal upon stimulation; the molecule, when experimentally applied, must produce a response in the postsynaptic cell that mimics the response produced by the release of neurotransmitter from the presynaptic neuron^69^. The neuroscience textbook most widely used internationally for the last 4 decades lists 4 criteria for a neurotransmitter ^70,71^: it is synthesized in the presynaptic neuron; it is present within vesicles and is released in amounts sufficient to exert a defined action on the postsynaptic neuron or effector organ; when administered exogenously in reasonable concentrations it mimics the action of the endogenous transmitter; a specific mechanism usually exists for removing the substance from the synaptic cleft. These are similar, but not identical, to the classic textbook on neurotransmitters: a neurotransmitter “should be synthesized and released presynaptically; it must mimic the action of the endogenous compound that is release on nerve stimulation; and where possible, a pharmacological identity is required where drugs that either potentiate or block postsynaptic responses to the endogenously released agent also act identically to the suspected neurotransmitter that is administered” ^222^. The pharmacological criterion is listed in another textbook ^223^.

Some authors note difficulties of establishing a CNS neurotransmitter. For example, a specialized neurotransmitter book states that “the candidate neurotransmitter should be present in the presynaptic terminal, be released when the presynaptic terminal is active, and when applied experimentally, induce faithful responses in the postsynaptic neuron. In practice, since central nervous system neurons continuously integrate diverse excitations and inhibitions, the last criterion is relaxed to demonstrating merely changes in such activity” ^224^.

Solomon Snyder, a leading scientist of classic neurotransmitters, neuropeptides and their receptors, wrote that “designating a molecule as a transmitter depends on the criteria employed, the most common of which are that the substance is synthesized in neurons, released by their terminals, mimics the effects of physiologic neurotransmission and possess a mechanism for inactivation. However, with each new candidate the rules have been modified and broadened” ^225^.

### Evidence supporting Cr as a neurotransmitter

16 small molecules have been listed as neurotransmitters in the classic textbook ^70,71^. Among them, adenosine, arachidonic acid, nitric oxide and carbon monoxide do not meet all four criteria, at the present. Cr appears to be better than these in meeting the criteria for a central neurotransmitter.

The results obtained by us in this paper have satisfied the criteria of Robinson ^224^ for Cr to be a CNS neurotransmitter.

The four criteria of Snyder and colleagues ^225^ have been mostly met but the physiological neurotransmission would require more research because a specific synapse(s) would have to be defined and studied for putative creatinergic neurotransmission. This can take much longer in the CNS than the PNS. Some commonly accepted neurotransmitters have never satisfied this criterion in a strict sense. The mechanism of Cr removal criterion is met not only by the Cr uptake in brain slices ^220^ and in synaptosomes ^221^, but also by our demonstration that SLC6A8 is required for synaptosome uptake of Cr.

The four criteria of Kandel et al.^70,71^ are mostly satisfied with some details requiring further research. The synthesis requirement is usually not strict because there are transmitters synthesized in some cells and transported into others where they function as transmitters. Our discovery of Cr in SVs can replace the synthesis requirement because the presence in neuronal SVs provide sufficient evidence that Cr is located in the right location to function as a neurotransmitter. The level of Cr in SVs is higher than those of ACh and 5-HT (Figs. 1 and 2). The amount of released Cr is in the same order of magnitude as those of Glu and GABA (Figs. 5 and 6). The criterion of a specific mechanism of removal was met by Cr uptake experiments in slices ^220^ and in synaptosomes ^221^, and further strengthened by our finding of SLC6A8 involvement in synaptosome uptake of Cr (Figure 8).

Here we report that Cr, at a concentration comparable to classical neurotransmitters, inhibits pyramidal neurons in specific regions of the mouse brain, with approximate 1/3 of pyramidal neurons responding to 100 μM Cr (Figure 7). In previous reports, 100 μM to 2 mM of GABA^226–228^, 50 μM to 2 mM of Glu ^229,230^, 1 to 100 μM of DA^229^, 0.1-100 μM of 5-HT^231–236^ were bath applied to investigate the physiological functions of neurotransmitters. Our results revealed that, when bath applied, Cr could inhibit cortical neurons at 100 μM within several minutes, with a time course similar to that of 5-HT^237 234 236^ and DA ^229^, but significantly slower than that of Glu^230^, GABA^48,227^ and 5-HT^233^.

In a recent report, knockout of the Slc6a8 gene increased excitation of cortical neurons^267^. Electrophysiological characterization of pyramidal neurons in the prefrontal cortex (PFC) found increased evoked firing frequency. Because we have shown that Cr inhibit a fraction of pyramidal neurons in the neocortex (Figure 8), this paper provides in vivo evidence consistent with the possibility of Cr as an inhibitory neurotransmitter.

### Differences between Cr and classic neurotransmitters

At this point, we do not have a molecularly defined receptor for Cr, only inferring its presence from the electrophysiological responses to Cr. We speculate that Cr may act on G-protein coupled receptors (GPCRs), rather than the fast-acting ligand-gated ion channels, such as AMPA or NMDA receptors for Glu and GABA_A_ receptor for GABA. There have been previous reports of Cr effects on neurons, including Cr as a partial agonist for GABA_A_ receptors ^238–241^. These effects require very high concentrations of Cr (in the ten millimolar range). There was also a report of the opposite effect: that Cr (at a concentration above 500 μM) increased neuronal excitability through NMDA receptors after incubation for 60 mins, with a time course significantly slower than those of classic neurotransmitters ^242^.

Ca^2+^ independent component of Cr release induced by extracellular K^+^ was more prominent than those of Glu or GABA. One possibility was that Ca^2+^-independent Cr release came from glia, because high GAMT levels were reported in astrocytes^261^ and oligodendrites ^261, 262^. As reported, other neuromodulators such as taurine can be released from astrocytes ^263^ or slices ^264^ in an Ca^2+^ independent manner. In addition, in the absence of potassium stimulation, Ca^2+^ depletion increased release of taurine in cultured astrocytes ^265^ or in striatum *in vivo*^266^. Similarly, in SLC6A8 KO slices, Ca^2+^ depletion (Figure 5G) also increased Cr baseline as compared to that in normal ACSF (Figure 5D).

With much longer history of research, ACh and 5-HT now have more evidence in other aspects than Cr as a central transmitter, especially because there are many agonists and antagonists for ACh and 5-HT to prove an additional criterion that is required in some ^222,223^, but not the majority of, textbooks for a neurotransmitter. The pharmacology criterion will take some time and efforts, because, so far, no efforts have been made to find agonists or antagonists for Cr.

### Implications of SLC6A8 and Cr

It is notable that SLC6A8 belongs to the neurotransmitter transporter family, with multiple members already shown to transport neurotransmitters ^65,66,90–99^.

The uptake experiments by others and us indicate that SLC6A8 transports Cr into neurons within the brain. AGAT is also expressed in the brain, but in cells not expressing SLC6A8 ^243,244^. Cr and its precursor were thought to be transported between different cells in the nervous system. When SLC6A8 was completely missing, such as in homozygous SLC6A8 deficient patients, Cr treatment was not effective. But, if

SLC6A8 was partially active, Cr was effective ^245^. Intractable epilepsy in a female with heterozygous SLC6A8 mutation was completely treated by Cr ^246^. Our data of inhibitory effect of creatine on cortical neurons might provide a new mechanism to its anti-epileptic activity ^247^.

The absence of SLC6A8 expression in astrocytes whose endfeet lining microcapillary endothelial cells (MCEC) form the blood-brain barrier (BBB) indicates that Cr in the brain does not rely on import from the periphery and is instead mainly synthesized in the brain ^80,82,248^. SLC6A8 functions within the brain to transport Cr and its precursors, not as a major contributor of Cr transport across the BBB. It is thought to mediate Cr uptake into the presynaptic terminal, based on studies of synaptosomes (Figure 8) ^221^.

Cr is known to have effects other than an energy source, and Cr supplement has been thought to be beneficial for children, pregnant and lactating women, and old people ^79,249^. Cr has been reported to improve human mental performance ^250–255^. Cr has been used as potential treatment in animal models of neurodegenerative diseases ^256,257^. Our work will stimulate further research to distinguish which of the previously suspected effects of Cr is not attributed to its role as an energy storage, but can be attributed to its role as a neurotransmitter.

### Search for new neurotransmitters

Our work may stimulate the search for more neurotransmitters. Our discovery indicates that the hunt for neurotransmitters stopped decades ago because of technical difficulties, not due to absence of more neurotransmitters. The fact that most of the known small molecule neurotransmitters have been found because of their peripheral effects also argues that what is missing is the concerted efforts to uncover central neurotransmitters with no peripheral effects. New neurotransmitters may be discovered from candidates which have been long suspected, and from previously unsuspected molecules or even previously unknown molecules.

Innovative approaches should be taken to uncover molecules with no previous suspicions or hints. Highly purified SVs, SVs from different regions of the brain, SVs with specific SLCs offer some of the starting points for future research.

## Materials and methods

Generation of knockout and knockin mice.

SLC6A8 knockout and knockin mice were generated using CRISPR-Cas9-mediated genome engineering techniques by Beijing Biocytogen (Beijing, China). AGAT ‘knockout-first’ ^185^ mice were purchased from CAM-SU GRC (Suzhou, China). All mutations were validated by southern blot analysis, tail junction PCR and DNA sequencing.

### Reverse-transcription polymerase chain reaction (RT-PCR) and quantitative real-time PCR (qPCR)

Total RNA of whole brains from mice of different genotypes was extracted using the Buffer RZ (Tiangen, no. RK14, Beijing, China) and reverse transcribed into complementary DNA (cDNA) using the RevertAid First-Strand cDNA synthesis kit (Thermo Scientific, K1622, USA). qPCR was performed using the Taq Pro Universal SYBR qPCR Master Mix (Vazyme, Q712-02) on Bio-Rad CFX-96 Touch Real-time PCR System (Bio-Rad, USA). Glyceraldehyde-3-phosphate dehydrogenase (GAPDH) was used as an internal control. ΔCt (difference in cycle threshold) was calculated for each sample (ΔCt = Ct _Target gene_ − Ct _GAPDH_) for further evaluation of relative mRNA expression levels in different genotypes. The sequence specificities of the primers were examined. Three pairs of primers targeting different genes were used: SLC6a8 forward, 5’-GTCTGGTGACGAGAAGAAGGG-3’, Slc6a8 reverse, 5’-CCACGCACGACATGATGAAGT −3’; AGAT forward, 5’-CACAGTGGAGGTGAAGGCCAATACATAT-3’, AGAT reverse, 5’-CCGCCTCACGGTCACTCCT-3’; GAPDH forward, 5’-AGGTCGGTGTGAACGGATTTG −3’, GAPDH reverse, 5’-TGTAGACCATGTAGTTGAGGTCA-3’.

Primers for reverse PCR were designed to obtain complete coding sequences based on information obtained from National Center for Biotechnology Information (NCBI): SLC6A8 forward, 5’-ATGGCGAAAAAGAGCGCTGAAAACG-3’; Slc6a8 reverse, 5’-TTACATGACACTCTCCACCACGACGACC-3’; AGAT forward, 5’-ATGCTACGGGTGCGGTGTCT-3’; AGAT reverse, 5’-TCAGTCAAAGTAGGACTGAAGGGTGCCT-3’. PCR products were electrophoresed on 1% agarose gels, stained with GelRed, visualized under UV illumination and photographed.

### Immunoblot analysis

Samples were loaded onto 10% polyacrylamide gels with the PAGE system (#1610183, Bio-Rad Laboratories, USA) and run in the SDS running buffer (25 mM Tris, 192 mM glycine, 0.1% SDS, pH 8.8) for 25 min at 80 V followed by 25 - 45 min at 200 V. Afterward, proteins were transferred to immobilon NC transfer membranes (HATF00010, Millipore) at 400 mA for 2 h in transfer buffer (25 mM Tris, 192 mM glycine, 20% methanol). Membranes were blocked in 5% fat-free milk powder in TBST (25 mM Tris, 150 mM NaCl, 0.2%Tween-20 (P1397, Sigma), pH 7.4 adjusted with HCl) and incubated overnight with the indicated primary antibodies dissolved in TBST containing 2% BSA.

Primary antibodies are listed below: rabbit anti-Synaptophysin (dilution 1:5000, cat no. 101002, SySy, Germany), rabbit anti-Synaptotagmin1/2 (dilution 1:2000, cat no.105002, SySy, Germany), rabbit anti-proton ATPase (dilution 1:1000, cat no.109002, SySy, Germany), rabbit anti-Synaptobrevin 2 (dilution 1:5000, cat no.104202, SySy, Germany), rabbit anti-SV2A (dilution 1:2000, cat no.109003, SySy, Germany), rabbit anti-VGlut1(dilution 1:4000, 135302, SySy, Germany), rabbit anti-VGlut2 (dilution 1:2000, 135402, SySy, Germany), rabbit anti-VGAT(dilution

1:4000, 131002, SySy, Germany), rabbit anti-SNAP23 (dilution 1:2000, cat no. 111202, SySy, Germany), mouse anti-PSD95 (dilution 1:5000, cat no 75028,NeuroMab), mouse anti-GluN1(dilution 1:5000, cat no 114011, SySy, Germany), rabbit anti-GM130 (dilution 1:1000, cat no ab52649, Abcam), rabbit anti-Golgin-97 (dilution 1:2000, cat no13192, Cell Signaling Technology, USA), rabbit anti-EEA1 (dilution 1:2000, cat no 3288, Cell Signaling Technology, USA), rabbit anti-LC3B (dilution 1:1000, cat no 2775S, Cell Signaling Technology, USA), goat anti-CathepsinB (dilution 1:2000, AF965, R&D Systems), rabbit anti-GAPDH (dilution 1:1000, cat no 2118S, Cell Signaling Technology, USA), rabbit anti-GluT4 (dilution 1:1000, ab33780, Abcam), rabbit anti-CACNA1A (dilution 1:300, 152103, SySy, Germany), rabbit anti-VDAC (dilution 1:1000, cat no 4661S, Cell Signaling Technology, USA), rabbit anti-MBP (1:1000, cat no. 295003, SySy, Germany), mouse anti Creatine Kinase B (dilution 1:5000, cat no MAB9076, R&D Systems), rabbit anti HA (dilution 1:2000, CST3724; Cell Signaling Technology, USA) and rabbit anti SNAP25 (dilution 1:2000, ab109105, Abcam) antibodies.

Membranes were washed in three washing steps in TBST (each for 5 mins) and incubated with peroxidase-conjugated secondary antibodies for 2-3 h at 4 °C. The second antibody used were anti-rabbit (dilution 1:5000, A6154, Sigma), anti-mouse (dilution 1:5000, 715-035-150, Jackson Immuno Research) or rabbit anti-goat IgG secondary antibodies (dilution 1:1000, cat no ab6741, Abcam). After repeated washing, signals were visualized using a ChemiDoc XRS^+^ System (Bio-RAD Laboratories, USA).

### Isolation of synaptic vesicles

Our purification procedures for synaptic vesicles were based on previously established immunoisolation methods ^143,151^. Protein G magnetic beads (cat no.88848, Thermofisher Scienctific) were washed three times with IP buffer (100 mM potassium tartrate, 4 mM HEPES-KOH, 2 mM MgCl_2_, pH 7.4) supplemented with a complete protease inhibitor cocktail (Roche). 5 μg monoclonal anti-Syp antibody directed against a cytoplasmic epitope (cat no 101011, synaptic systems, Germany) or control mouse IgG (10400C, Thermofisher Scientific) was used to incubate with 20-30 μl beads for 30 mins at RT in 2% BSA dissolved in IP buffer. Under this condition, 4-4.5 μg of antibody was coupled, as determined by Western blot and Commassie Blue staining. Immunoisolation of SVs were carried out at 0 °C-2 °C to prevent vesicular content leakage (with RT as a control). Briefly, the whole mouse brain was homogenized in 3 ml of IP buffer with a glass/Teflon homogenizer (20 strokes at 2000 rpm, WHEATON, USA and WIGGENS WB2000-M, Germany) immediately after decapitation. Homogenates were centrifuged for 25 mins at 35,000 x g and the supernatant was adjusted to approximately 3 mg/ml protein (Nanodrop 2000 C, Thermofisher Scientific). To capture the SVs for content detection, about 200 μl of supernatants (per 5 μg anti-Syp/IgG) were incubated with pre-coupled beads for 2.25 h under slow rotation at 2 °C. Beads were washed 6 times for further Western blot analysis and vesicular content detection. For pharmacological blockade of H^+^-gradient across SV membrane, the mix of supernatants and pre-coupled beads was diluted into 1.2 ml before the addition of inhibitors.

### Determination of vesicular contents

To extract SV contents, immunoisolates were treated with 50 μl ultra-pure water. 100 μl methanol together with 100 μl acetonitrile were added to precipitate proteins in samples. After centrifugation for 20 mins at 168000 × g, supernatants were collected and centrifuged for 20 mins at 2000 × g to remove beads and proteins. Samples were pre-frozen with liquid nitrogen and vacuum dried at −45 °C overnight. Dried samples were kept frozen and re-suspended with 50 μl of 0.2 μM ^13^C-creatine (internal control) immediately before detection.

Capillary electrophoresis-mass spectrometry (CE-MS) was used to verify and quantify small molecules. CE/MS detection was applied with the coupling of PA800 plus CE system (Beckman Coulter, Brea, CA, USA) and mass spectrometry (TRIPLE QUAD 5500, AB SCIEX or Q Exactive HF-X, Thermo Scientific). Before SV content detection, we optimized MS detection of classical neuro transmitters, Cr and amino acids in positive ion mode. Firstly, the fragment ions (Q3) for a given molecule (precursor ions, Q1) were determined by either systematic scanning of standard sample solution (0.1 μM in 10% acetate acid) or referring to database (www.mzcloud.org). Secondly, optimal values of collision energy (CE), collision cell exit potential (CXP) and declustering potentials (DP) were determined for each pair of Q1/Q3. Thirdly, optimal combination of parameters (Q1/Q3, CE, CXP, DP) was chosen for each molecule. In addition, parameters were adjusted every 2 to 3 months for best signal to noise ratios.

CE/MS separations were carried out by capillaries (OptiMS silica surface cartridge, BECKMAN COULTER). The CE background electrolyte was 10 % acetate acid. Each new separation capillary was activated with rinsing under100 psi sequentially with methanol for 10 mins (forward), methanol for 3 mins (reverse), H_2_O for 10 mins (forward), H_2_O for 3 mins (reverse), 0.1 M NaOH for 10 mins (forward), water for 5 mins (reverse), 0.1 M HCl for 10 mins (forward), followed by water for 10 mins and then 10 % acetate acid for 10 mins (forward) and 3 mins (reverse), prior to the first use. Between analyses, the capillary was rinsed with 10% acetate acid under a 100 psi pressure for 5 min (forward) flowed by 75 psi for 4 min. The sample (50 μl) was injected with 2.5-4 psi for 30 s. Separation voltage of 25 kV was applied for 25 mins. To maintain stably spray during CE separation, ion spray voltage was applied at 1.7-1.9 kV. MS data were collected 5 mins after CE separation. Finally, the capillary was washed with 10% acetate acid for 10 mins, followed by methanol for 20 min and then10% acetate acid for 20 mins.

Standard solutions of 0.2 μM ^13^C-Cr (internal control) and analytes were used to plot standard curves. Linear standard curves (R^2^>0.98, for most cases, R^2^>0.99), calculated from peak area ratios corresponding to analytes and internal standards, were obtained for all molecules tested. The concentration ranges used for standards of Glu, GABA, ACh, 5-HT, Cr and alanine were 0.03-10 μM, 0.003-1 μM, 0.0003-0.1 μM, 0.003-1 μM, 0.03-1 μM and 0.03-1 μM, respectively. Standard curves were made at least twice for a given capillary. Analytes of SV contents were calculated using the standard curves and normalized to the amount of anti-Syp antibody conjugated to the beads.

### Electron microscopy

All EM grids were glow discharged for 30 s using a plasma cleaner (Harrick PDC-32G-2). To free SVs from beads, 25 μl 0.1 M glycine-HCl (PH=2) was incubated for 1 min and quickly neutralized with 25 μl 0.1 M Tris (pH=10). Beads were quickly removed and 2-4 μL aliquots of SVs were applied to the carbon coated copper grids (Zhong Jing Ke Yi, Beijing). After 1 min, the grid was dried with a filter paper (Whatman No.1), and placed in the water, and then immediately stained using 2 % uranyl acetate for 30 s. At last uranyl acetate was removed and the grid was air dried. The grids were examined on a JEM-F200 electron microscope (JEOL, Japan) operated at 200 kV. Images were recorded using a 4k × 4k COMS One view camera (Gatan). Fixation of synaptosomal pellets was performed by immersion with pre-warmed 2.5% glutaraldehyde in 0.1 M phosphate buffer (pH 7.4) at RT for 2 h. After washing 4 times with 0.1 M phosphate buffer (pH 7.4) every 15 mins, samples were post-fixed with 1% osmium tetroxide (w/v) at 4°C for 1 h and then washed 3 times. Following en bloc staining with 2% uranyl acetate (w/v) at 4°C overnight, samples were dehydrated and embedded in fresh resin, polymerized at 65°C for 24 h. Ultrathin (70 nm) sections were obtained by Leica UC7 ultramicrotome (Leica Microsystems, German) and recorded on 80 kV in a JEOL Jem-1400 transmission electron-microscope (JEOL, Japan) using a CMOS camera (XAROSA, EMSIS)

### Immunohistochemistry

Adult mice were anesthetized by *i.p.* injection with 2 % 2,2,2-tribromoethanol (T48402, Sigma Aldrich) in saline at a dose of 400 mg/kg, and perfused trancardially with 0.9 % saline followed by 4 % PFA in PBS (137 mM NaCl, 2.7 mM KCl, 10 mM Na_2_HPO_4_, 1.8 mM KH_2_PO_4_, pH=7.4).

Brains were cryoprotected with 30 % sucrose in 30 % sucrose 0.1 M PB (81 mM Na_2_HPO_4,_ 19 mM NaH_2_PO_4_) and sectioned in the coronal plane (40 μm thick) using a Cryostat (Leica 3050S). For anti-HA immunostaining, we used a rabbit monoclonal anti-HA antibody (1:500 in 0.3 % Triton in PBS; 48 h incubation at 4 °C; #3724, Cell Signaling Technology), followed by a goat anti-rabbit Alexa Fluor 546 secondary antibody (1:1000; overnight at 4 °C; # A-11035, Invitrogen). Sections were mounted in a medium containing 50 % glycerol, cover-slipped, and sealed with nail polish. Images were acquired using virtual slide microscope (Olympus VS120-S6-W) and a laser-scanning confocal microscope (Zesis 710) and brain structures inferred with an established mouse brain atlas ^258^.

### Preparations of brain slices

Male C57 mice (of 30-38 days-old) were anesthetized with pentobarbital (250 mg/kg) and decapitated. Brains were quickly removed and placed into ice-cold, low calcium, high magnesium artificial cerebrospinal fluid (ACSF) with sodium replaced by choline.

The medium consisted of 120 mM choline chloride, 2.5 mM KCl, 7 mM MgSO_4_, 0.5 mM CaCl_2_, 1.25 mM NaH_2_PO_4_, 5 mM sodium ascorbate, 3 mM sodium pyruvate, 26 mM NaHCO_3_, and 25 mM D-(+)-glucose, and was pre-equilibrated with 95% O2-5% CO2. Coronal brain slices (300 μm thick) were cut with a vibratome (Leica VT1200S). Slices were incubated for 1 h at 34°C with oxygenated ACSF containing: 124 mM NaCl, 2.5 mM KCl, 2 mM MgSO_4_, 2.5 mM CaCl_2_, 1.25 mM NaH_2_PO_4_, 26 mM NaHCO_3_, and 10 mM D-(+)-glucose.

### Evoked release from brain slices

Coronal brain slices (each 300 μm thick, typically with a wet weight of 17-20 mg) were transferred into a specially designed superfusion chamber with a volume of approximately 200 μl, containing freshly 95% O_2_/5% CO_2_ oxygenated ACSF. Slices were equilibrated for 10 min in ACSF at a superfusion rate of 0.9-1.25 ml/min. The “control” sample was collected for 1 min just before high K^+^ stimulation (K-ACSF, 70 mM KCl replacing equal amount of NaCl). We waited for 30 seconds to allow K^+^ stimulus to immerse the slices (dead volume for solution transition of 200 μl and chamber volume of 200 μl), then the sample “70 mM K” in response to K-ACSF was collected for another 1 min. Following 10 min of washout period, we collected the third sample of “wash” for 1 min.

To detect Ca^2+^ dependent release, slices were pre-incubated for 10 mins with normal ACSF and equilibrated with Ca^2+^ free ACSF (containing 1 mM EGTA to chelate extracellular Ca^2+^) for 10 mins. The baseline sample “0 Ca^2+^ ACSF” was collected for 1 min. Superfusion solution was changed to Ca^2+^ free K-ACSF for 2 mins and sample “0 Ca^2+^ 70 mM K” was collected (dead volume for solution transition of 400 μl and chamber volume of 200 μl). After that, the solution was changed back to normal ACSF for 10 mins and K-ACSF for 2 mins. The sample “2.5 mM Ca 70 mM K” for the last min was collected.

Samples were subjected to CE-MS in a method similar to SV content detection, except for the following: 1) standards were dissolved in ACSF or other buffers used in release experiment; 2) concentration ranges used for standards of Glu was from 0.003-1 μM. 3) To protect the MS from salt pollution, data were collected from 10 mins −20 mins during CE separation.

### Patch-Clamp recordings

Slices were transferred to a recording chamber on an upright fluorescent microscope equipped with differential interference contrast optics (DIC; Olympus BX51WI). Slices were submerged and superfused with ACSF at about 2.8 ml/min at 24-26 °C. Whole cell patch recordings were routinely achieved from layer 4/5 medium sized pyramidal neurons from the somatosensory cortex. Patch pipettes (3-5 MΩ) contained: 140 mM K-gluconate, 10 mM HEPES, 0.5 mM EGTA, 5 mM KCl, 3 mM Na_2_-ATP, 0.5 mM Na_3_GTP, and 4 mM MgCl_2_ (with pH adjusted to 7.3 and osmolarity of 290 mOsm/kg). Current-clamp recordings were carried out with a computer-controlled amplifier (Multiclamp 700B, Molecular Devices) and traces were digitized at 10 kHz (DigiData 1550B, Molecular Devices). Data were collected and analyzed using Clampfitor Clampex 10 software (Molecular Devices).

Cells were characterized by their membrane responses and firing patterns during hyperpolarizing and depolarizing current steps (−100 to +500 pA, increment: 50 pA or 25 pA, 500 ms). Regular spiking pyramidal neurons were identified by moderate maximal spiking frequencies (20-60 Hz, that is, 10-30 spikes per 500 ms, Figure 7E-G), increasing of inter-spike intervals during depolarizing step (Figure 7B and Supplementary Figure 11A), high action amplitude (Supplementary Figure 12E) and large half width (Supplementary Figure 12F) ^201,202^. After the mean firing frequency evoked by current injections reached the steady state for at least 5 mins (typically 20-30 mins following the formation of whole-cell configuration), 100 μM Cr was bath-applied for 6 mins. Typically, Cr was applied for a 2^nd^ time following washout to re-confirm the effects.

### Synaptosome preparation

Synaptosomes were isolated by Ficoll/sucrose density-gradient centrifugation ^203,209,221,259^. Whole brains from adult male mice were homogenized with 15 strokes at 900 rpm in buffer A (320 mM sucrose, 1 mM EDTA, 1mM EGTA, 10 mM Tris–HCl, pH 7.4, with a complete protease inhibitor cocktail (Roche)). The homogenate (H fraction) was centrifuged at 1000 g for 10 min to precipitate the membrane fragments and nuclei (P1 fraction). Supernatant was centrifuged again at 1000 g for 10 min and the resulting supernatant (S1) was centrifuged at 12,000 g for 20 min. Supernatant was the S2 fraction, and the pellet was re-suspended with buffer A and centrifuged at 12,000 g for 20 min. The resulting pellet was crude synaptosomes (P2 fraction), containing synaptosomes with mitochondria and microsomes.

Crude synaptosomes (P2 fraction) was re-suspended with 150-200 μl buffer B (320 mM sucrose and 10 mM Tris–HCI (pH 7.4)). The sample was carefully overlaid on the top of a gradient of 2 ml of 7.5 % (wt/vol in buffer B) Ficoll and 1.8 ml of 13% (wt/vol in buffer B) Ficoll and centrifuged at 98,000 g for 45 min at 2-4 °C in a swinging-bucket rotor. A myelin band was present near the surface, and th synaptosomes band (fraction Sy) was present at the interface between the 13% and 7.5 % Ficoll layers, with the mitochondria being pelleted at the bottom. For further western analysis, the supernatant was divided into 6 fractions (600 μl for each fraction) and the mitochondria pellet was discarded. The isolated synaptosomes was included in fraction 4.

For Western analysis, fractions H, S1, P1, S2, P2 and Sy were adjusted to 0.5 mg/ml by bicinchoninic acid assay (BCA) method with reference to NanoDrop™ 2000 Spectrophotometers. 3.35 μg protein was loaded for each lane. Fractions 1-6 were loaded with the same volume (10 μl composed of 6.7 μl sample and 3.3 μl loading buffer) for each lane.

### Creatine uptake into synaptosomes

To remove Ficoll, we diluted the synaptosomal band (480 μl) with 4.3 ml of a pH 7.4 buffer C containing (in mM) 240 mannitol, 10 glucose, 4.8 Kgluconate, 2.2 Cagluconate, 1.2 MgSO4, 1.2 KH_2_PO4 and 25 HEPES-Tris. The sample was then centrifuged at 12,000 g and the pellet was re-suspended with buffer C. Uptake experiments were either performed at 37°C or at 0°C (control). For each sample, 25-43 μg of synaptosomes (with a volume of 40-50 μl) were added to 360 μl buffer containing (in mmol/L) 100 NaCl, 40 mannitol, 10 glucose, 4.8 Kgluconate, 2.2 Cagluconate, 1.2 MgSO_4_, 1.2 KH2PO_4_, 25 HEPES and 25 Tris (pH adjusted to 7.4). A mixture of 18 μM [^14^C]-creatine (0.4 μCi) and 5 μM creatine was quickly added. After 10 min, uptake was terminated by the addition of 1 ml of NaCl-free ice-cold buffer C. Samples were immediately filtered, under vacuum, through a Whatman GF/C glass filter (1825-025) pre-wetted with buffer C. Filters were further washed with 10 ml of ice-cold buffer C, dissolved in scintillation fluid and the radioactivity determined by liquid scintillation spectrometry.

### Creatine uptake into SVs

The uptake of ^13^C-creatine was assayed according to a conventional procedure ^260^ with slight modifications: the immunoisolated synaptic vesicles by 10 μg Syp antibody (101011, Synaptic Systems, Germany) were resuspended with the uptake buffer (150mM meglumine-tartrate, 4 mM KCl, 4 mM MgSO_4_, 10mM HEPES-KOH pH 7.4, and complete EDTA-free protease inhibitor cocktail) containing 4mM Mg-ATP or additional 4mM MgSO4, followed by preincubation for 30 min at 25L. The uptake reaction was started by addition of 1mM ^13^C-creatine dissolved in the uptake buffer with a final volume of 125 μL (pH at 6.8). After 10min at 25L, 1mL of ice-cold uptake buffer was added to the incubation to stop the reaction, followed by 5 more times washing. The SV contents were extracted using the protocol described in the determination of vesicular contents part. 100 nM Cr was used as the internal control. CE-MS and LC-MS was used to verify and quantify the creatine contents of samples. A Vanquish UHPLC system coupled to a Q Exactive HF-X mass spectrometer (both instrument Thermo Fisher Scientific, USA) were used for LC-MS analysis along with SeQuant ZIC-HILIC column (150 mm × 2.1 mm, 3.5 μm, Merck Millipore, 150442) in the positive mode and SeQuant ZIC-pHILIC column (150 mm × 2.1 mm, 5 μm, Merck Millipore, 150460) in the negative mode. For ZIC-HILIC column, the mobile phase A was 0.1% formic acid in water and the mobile phase B was 0.1% formic acid in acetonitrile. The linear gradient was as following: 0 min, 80% B; 6 min, 50% B; 13 min, 50% B; 14 min, 20% B; 18 min, 20% B; 18.5 min, 80% B; 30 min, 80% B. The flow rate used was 300 μl/min and the column temperature was maintained at 30°C. For ZIC-pHILIC column, the mobile phase A is 20 mM ammonium carbonate in water, adjusted to pH 9.0 with 0.1% ammonium hydroxide solution (25%) and the mobile phase B is 100% acetonitrile. The linear gradient was as follows: 0 min, 80% B; 2 min, 80% B; 19 min, 20% B; 20 min, 80% B; 30 min, 80% B. The flow rate used is 150 μl/min and the column temperature is 25°C. Samples were maintained at 4°C in Vanquish autosampler. 3 µl of extracted metabolites were injected for each run. IP samples were subjected to ZIC-HILIC column in positive mode for major metabolites detection, and then subject to ZIC-pHILIC column in negative mode for orthogonal detection.

## Acknowledgements

We are grateful to Drs. Xiaohui Zhang, Xinxiang Zhang, Minmin Luo, Qingchun Guo, Wuping Ge for discussion, suggestions and technical support, to Jiang Chen, Jing Cai, Yulin Jiang, Jinhuan Ou, Xiaomeng Deng, Jiawen Liu and Meng Wu for technical assistance, to CIBR, Peking-Tsinghua Center for Life Sciences, Changping Laboratory, Chinese Academy of Medical Sciences (2019RU003) and National Natural Science Foundation of China (81473189) for support. Research in the Rao lab has never been contaminated by the Chinese Brain Initiative.

## Author contributions

YR conceived the idea of searching for new neurotransmitters and supervised the project. YR, JZ, XJ and XB designed the experiments, JZ and XJ tried different approaches to find new neurotransmitters. XB improved the methods, carried out experiments and supervised technicians to carry out the experiments for this paper. XB, JZ, XJ and SY reproducibly found Cr in SVs in independent experiments. WL carried out the uptake experiments. ZL carried out experiments. YR, XB, XJ and JZ wrote the manuscript.

## Supplementary materials

**Supplementary Figure 1. Validation of SV purification from the mouse brain.** (A) A diagram for SV purification and SV content analysis. Brain homogenates were centrifugated at 35000 g for 25 mins and supernatants containing SVs were collected as the starting material. Following immunopurification by the anti-Syn antibody, SVs were broken by water to release their contents. Metabolites were analyzed by CE-MS. (B) EM of SVs captured by the anti-Syn antibody. Arrows pointing to SVs, the size each being 40.4± 0.26 nm, with size distribution shown on the right. (C) Analysis of SV purity by immunoblotting. Markers for SVs (Syp, syt, H^+^-ATPase, syb2, SV2A, VGLU1, VGLU2 and VGAT) were effectively pulled down by beads coated with the anti-Syn antibody but not by those coated with IgG. Non-SV markers present in supernatants, including those for the lysosome (LAMP1, cathepasinB, LC3B), the Golgi apparatus (GM130, Golgi 97), the early endosome (EEA1), the cytoplasma (GAPDH), synaptic membrane (SNAP23) and postsynaptic components (PSD95, GluN1), the mitochondria (VDAC), the cytoplasmic membrane (CACNA1A), axonal membrane (GluT4), and glia membrane (MBP), could not be pulled down by either the anti-Syp antibody or IgG.

**Supplementary Figure 2. Effects of pharmacological inhibitors on SV contents.** (A) Cr (B) Glu (C) GABA (D) ACh (E) 5-HT (F) Alanine*, p<0.5, **p<0.05, ***, p<0.01, one-way ANOVA with Tukey’s correction (n=4 per group). Data were normalized to average amount of molecules pulled down by the anti-Syp antibody. NIG, nigrecin.

**Supplementary Figure 3. Validation of SLC6A8 knockout mice.** (A) RT-PCR analysis of full-length coding sequences showing absence of SLC6A8 mRNA expression in brains from hemizygous (SLC6A8^-/Y^) male knockout mice. GAPDH serves as the control. (B) Detection of SLC6A8 mRNA expression in female mice, showing reduced expression in heterozygous (SLC6A8^+/-^) and loss of SLC6A8 expression in homozygous (SLC6A8^-/-^) knockout mice. (C and D) Quantitative RT-PCR analysis. The average values of SLC6A8 mRNA/GAPDH in WT mice (SLC6a8^+/Y^ males in D and SLC6A8*^+/+^* females in **e**) were set to 1. **, p<0.01 in (D), student t test. *** in (E), p<0.001, one-way ANOVA with Tukey’s correction (n=4 mice per group, with 3 repeats for each mice).

**Supplementary Figure 4. Brain and body weights of SLC6a8 knockout mice.** (A) Brain weights of male mice. (B) Body weights of male mice. (C) Brain weight of female mice. (D) Body weights of female mice. **, p<0.01 in (B), student t test. *, p<0.05, **, p<0.01,*** in (**e**), p<0.001, one-way ANOVA with Tukey’s correction (n=5 mice per group, 7 weeks old).

**Supplementary Figure 5. Representative CE-MS data of molecules.** (A-E) Molecules pull-down from WT mice by IgG (WT-IgG), anti-Syp antibody (WT-Syp), and those from knockout mice (SLC6A8KO-IgG, SLC6A8 KO-Syp). (A) Glu. (B) GABA. (C) ACh. (D) 5-HT. (E) Alanine. Vertical: signal amplitude; abscissa: retention time.

**Supplementary Figure 6. Proteins and small molecules detected in SVs from WT and SLC6a8 KO mice.** (A) Immunoblotting analysis of SV markers Syp, syt, H-ATPase and postsynaptic components (PSD95). (B-D) Peak areas of CE-MS signals for neurotransmitters: Glu, GABA, and ACh. These neurotransmitters are enriched in SVs and not affected by SLC6A8 genotypes. (E) Alanine was not enriched in SVs of either WT or SLC6A8 KO mice. (F) Representative raw example traces of Cr. (G) Cr in SVs was lower in SLC6A8 KO mice than that in WT mice. **, p<0.01, ***, p<0.001, one-way ANOVA with Tukey’s correction (n=8 samples per group).

**Supplementary Figure 7. Validation AGAT-KO first mice.** (A) RT-PCR analysis of the entire coding region showing the absence of AGAT mRNA in AGAT^-/-^ mice and decreased expression in AGAT^+/-^ mice. (B) Quantitative RT-PCR analysis. ***, p<0.001, one-way ANOVA with Tukey’s correction (n=4 mice per group, with 3 repeats for each mice).

**Supplementary Figure 8. Brain and body weights of *Agat* ^+/+^, *Agat* ^+/-^ and *Agat* ^-/-^ mice.** (A) Brain weights of male mice. (B) Body weights of male mice. *, p<0.05, **, p<0.01, * in (E), p<0.001, one-way ANOVA with Tukey’s correction, ns, not significant. (n=5 mice per group, 7-weeks).

**Supplementary Figure 9. Similar amounts of SVs pulled down from Agat +/+, Agat +/- and Agat -/- mice.** (A) Representative Western analysis of SV markers H+-ATPase, Syt and Syp from input, samples pulled down by IgG (IgG-P) and anti-Syp antibody (Syp-P). (B-E) Quantification of protein band densities for Syp (B), Syt (C) and H+-ATPase (D) (n=20 per group).

**Supplementary Figure 10. Layer distribution of recording sites in the somatosensory cortex.** (A) High levels of SLC6A-HA signal observed in the somatosensory cortex, especially in layer 4. (B) Recording sites. Red denotes the responsive neurons and blue the unresponsive neurons.

**Supplementary Figure 11. Electrophysiological parameters of Cr-responsive (blue) and Cr-nonresponsive neurons (red).** There was no significant difference between the two groups of neurons in membrane capacitance (Cm) (A) and membrane resistance (Rm) (B), resting membrane potential (RMP) (C), spike threshold (D), spike amplitude (E), half width of spike (F) and amplitude of after hyperpolarization (AHP) (G). While no electrical parameters detected were changed by Cr in non-responsive neurons (n=35), resting membrane potential was slightly inhibited by Cr and AHP was slightly elevated (n= 16) (*, p<0.05, paired t test).

**Supplementary Figure 12. Electrophysiological data of Cr-unresponsive neurons.** (A) Representative raw traces. (B) I-V curve of this same neuron. No effect of 100 μM Cr on rheobase current and evoked spike number at a current of 50 pA above rheobase current was observed.

