## Supplementary figures and images for "Evidence suggesting creatine as a new central neurotransmitter: presence in synaptic vesicles, release upon stimulation, effects on cortical neurons and uptake into synaptosomes and synaptic vesicles"

### FIgure S1

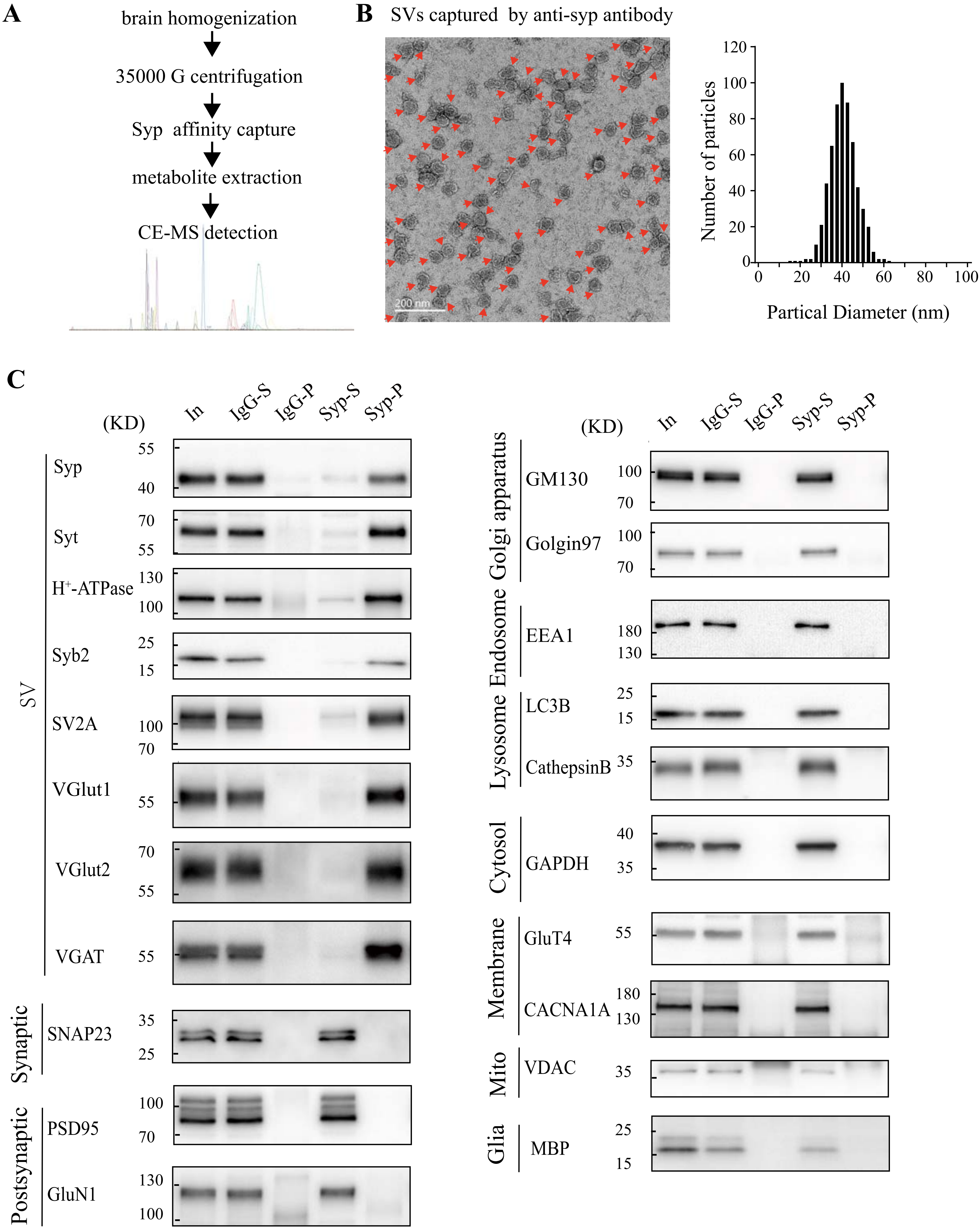

### FIgure S2

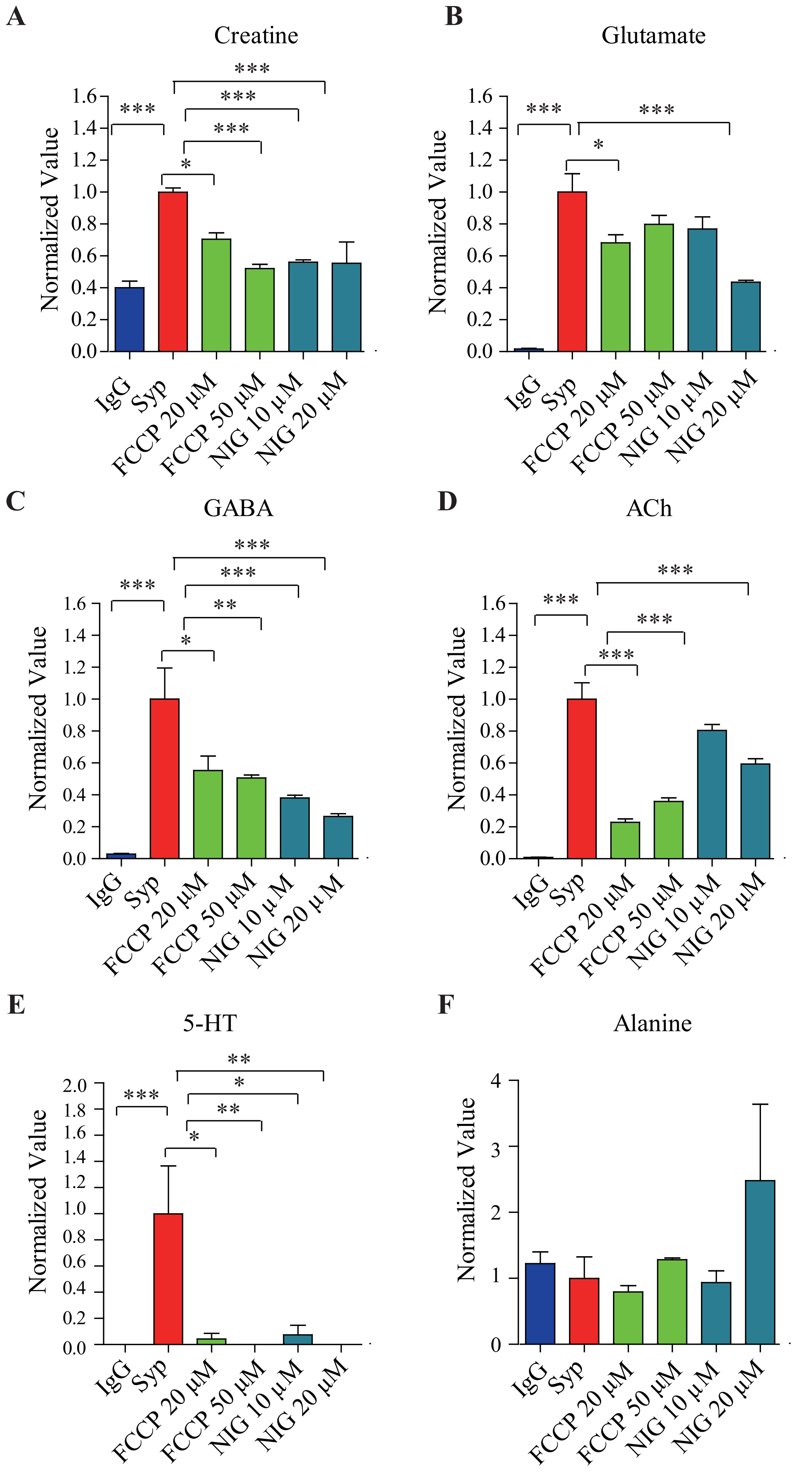

### FIgure S3

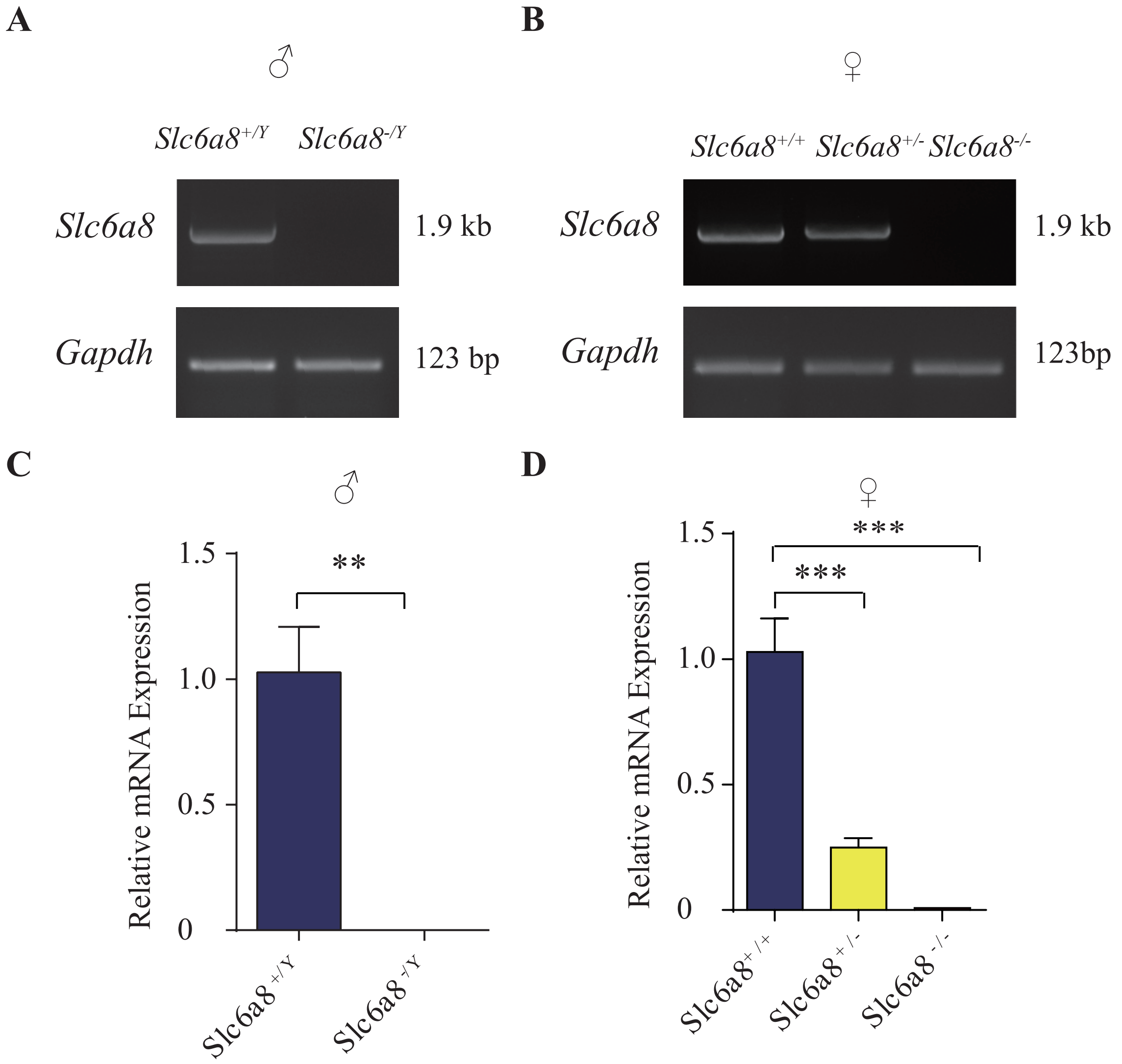

### FIgure S4

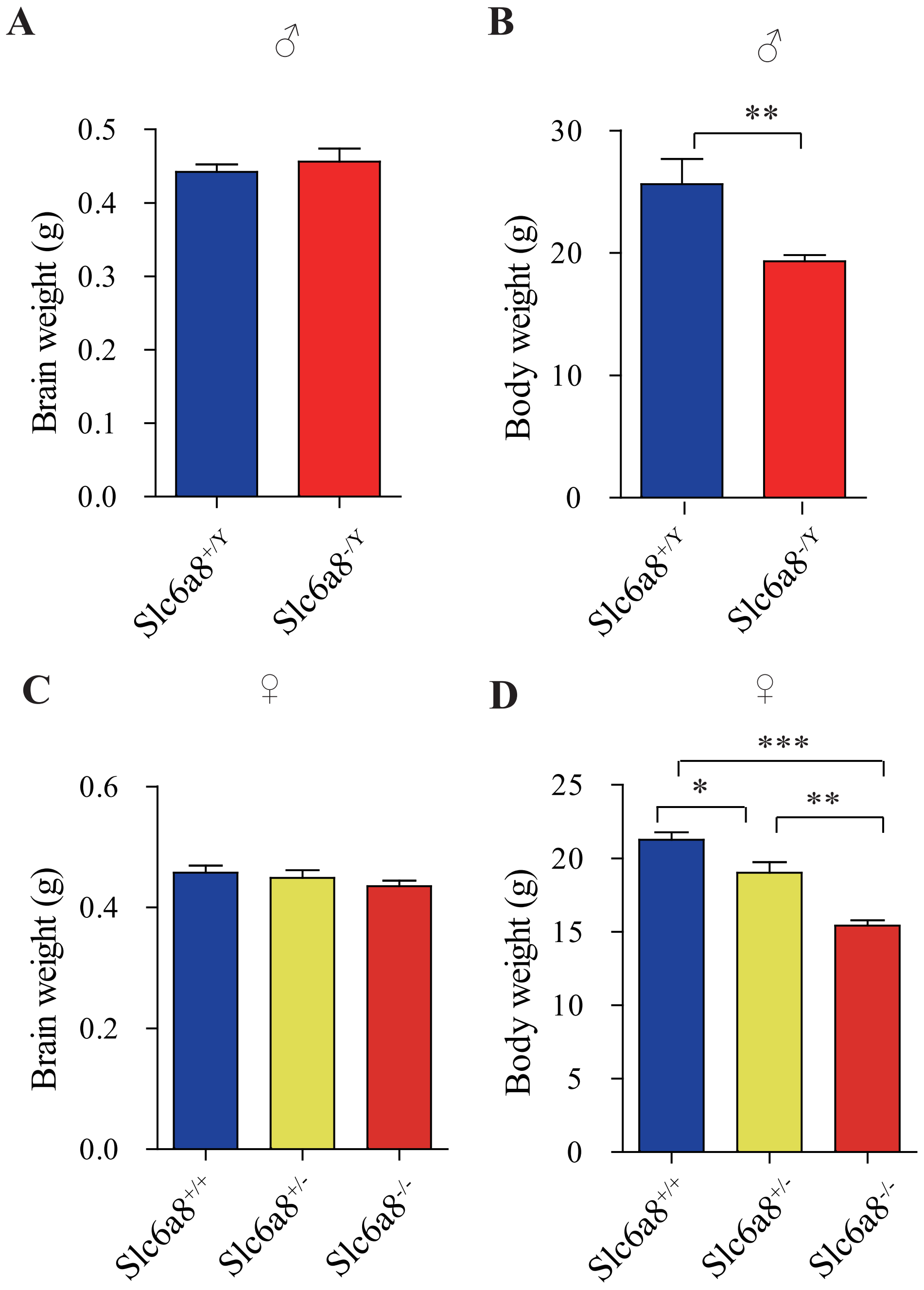

### FIgure S5

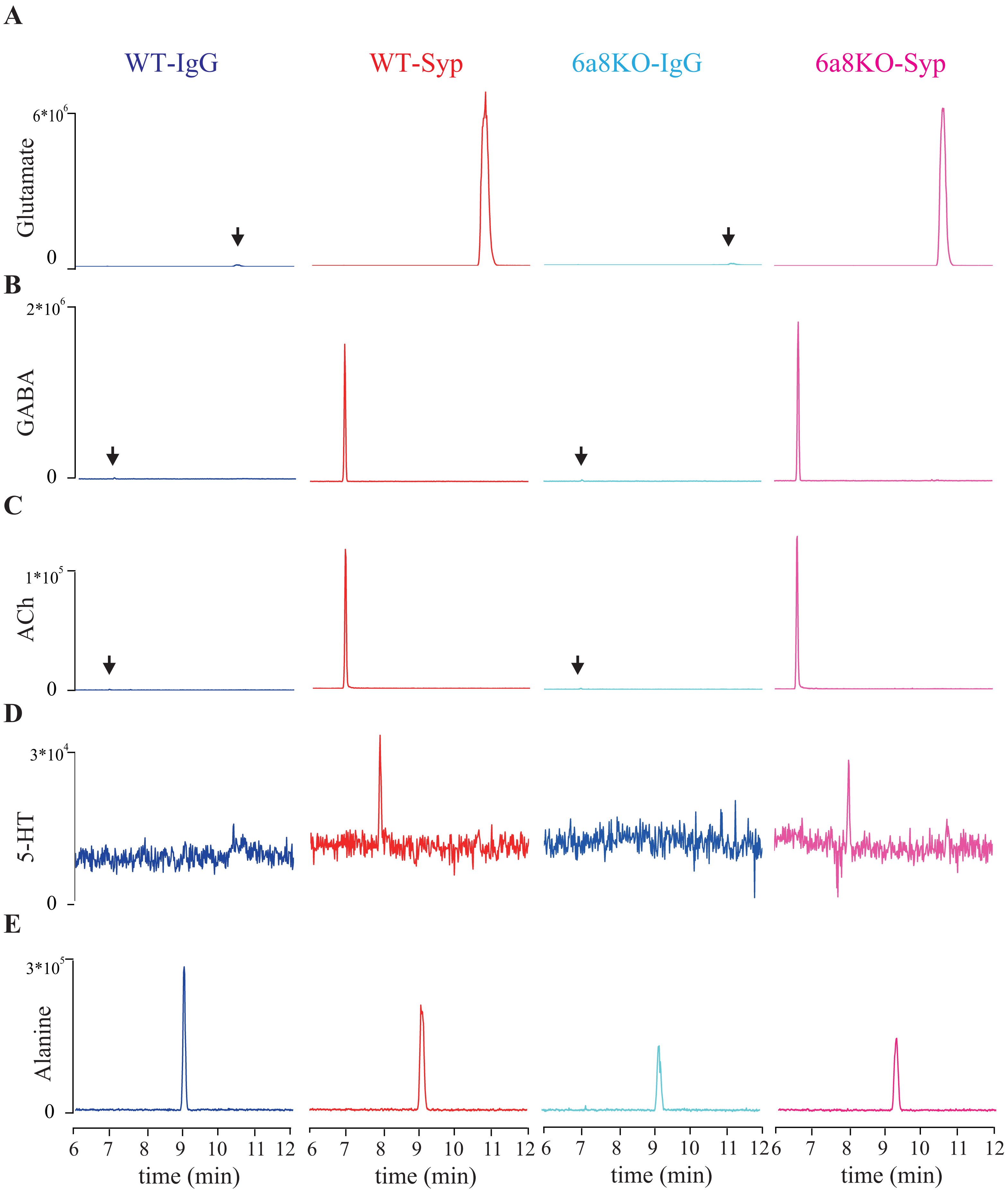

### FIgure S6

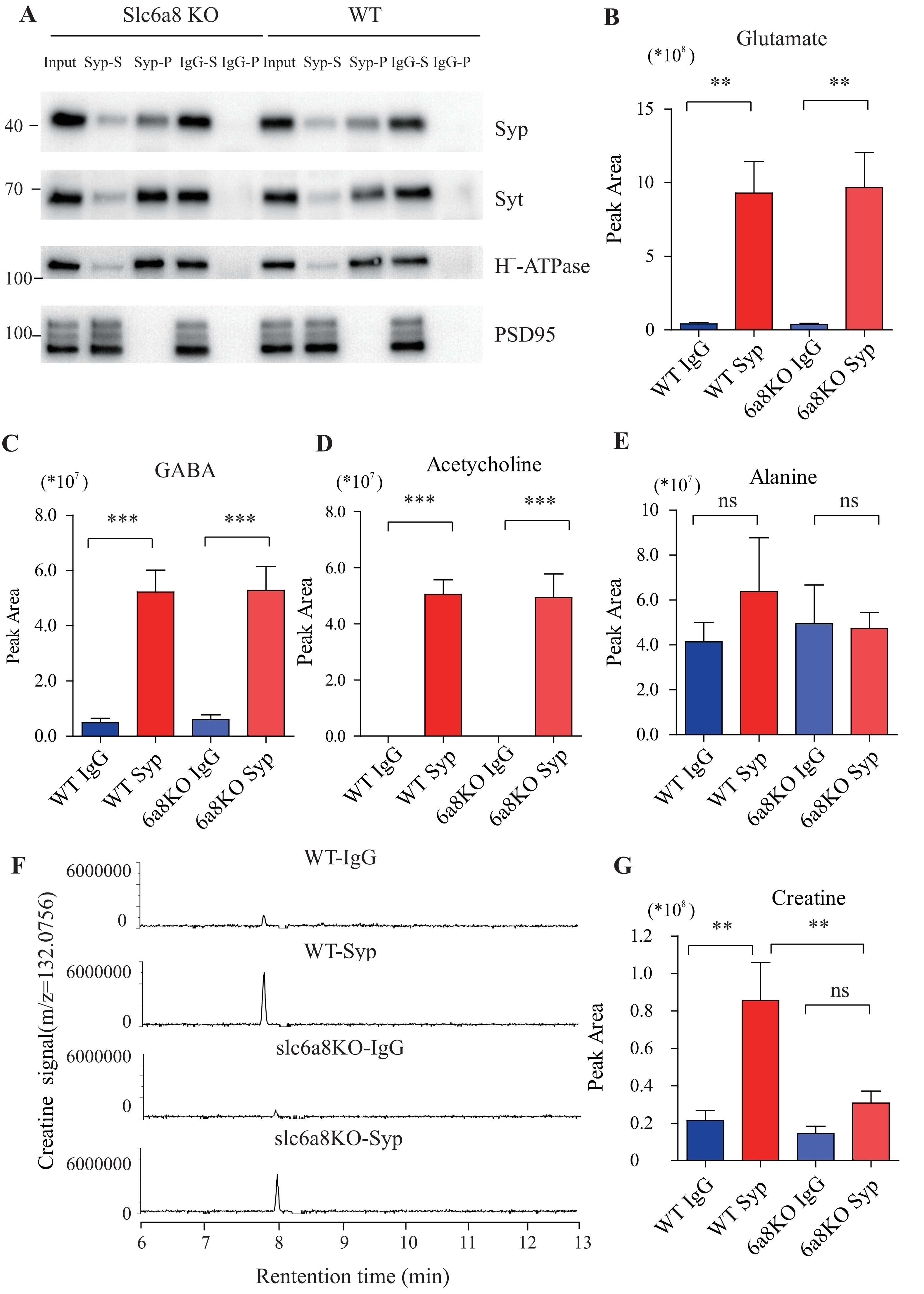

### FIgure S7

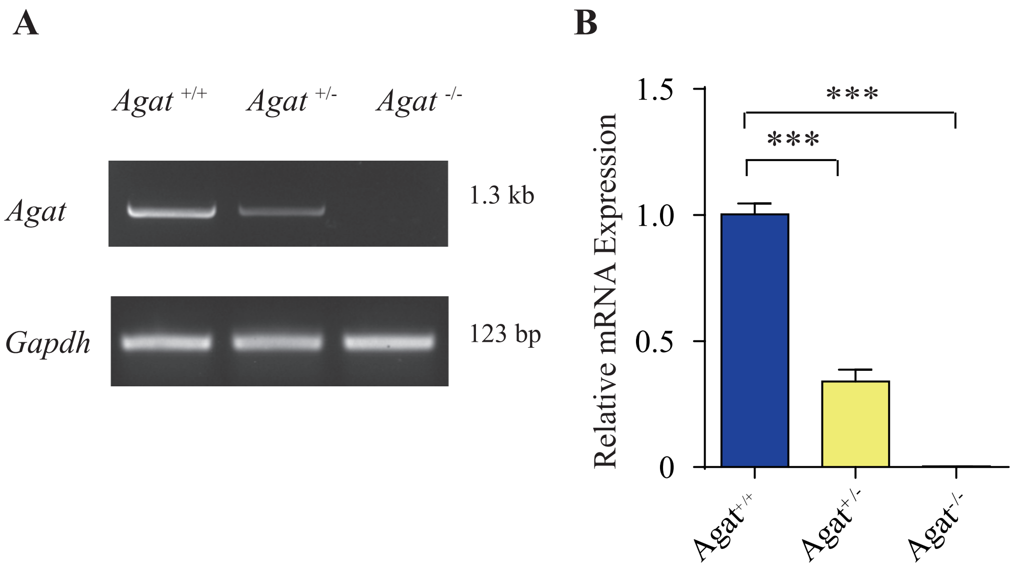

### FIgure S8

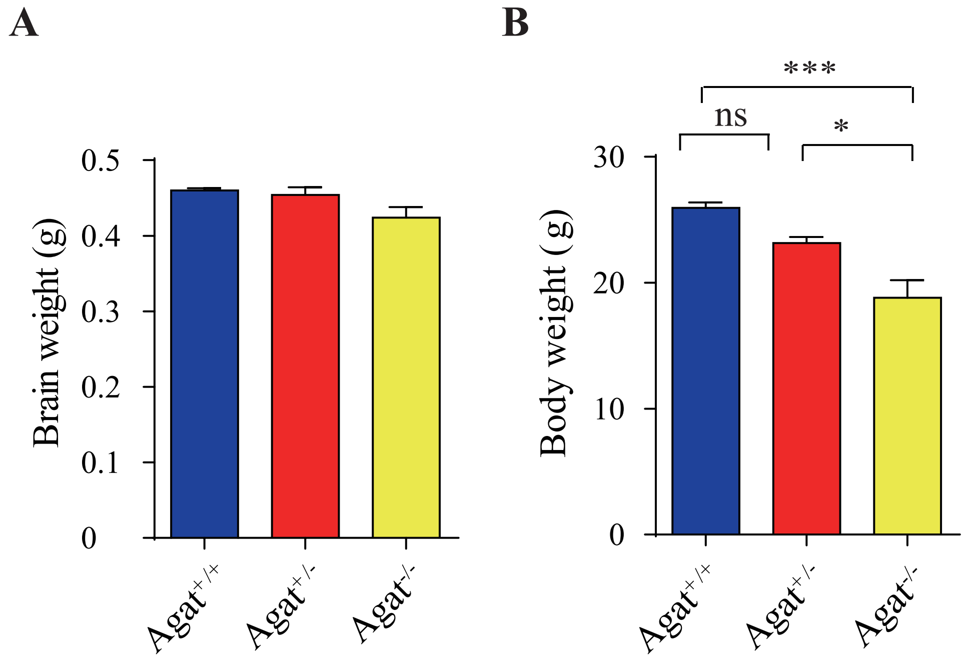

### FIgure S9

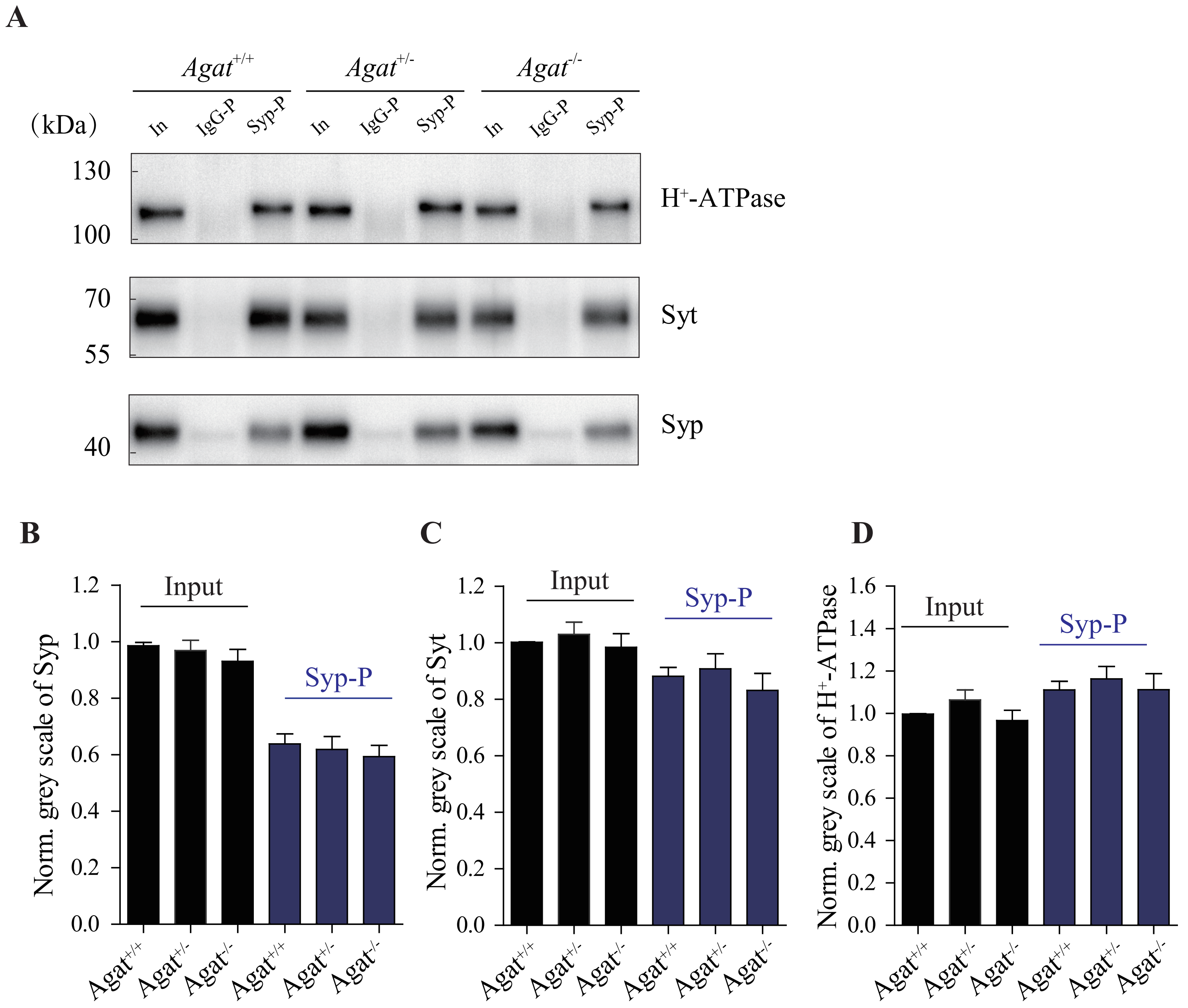

### FIgure S10

A

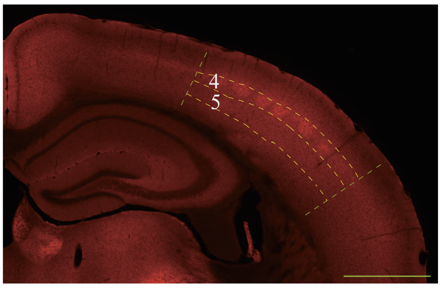


B

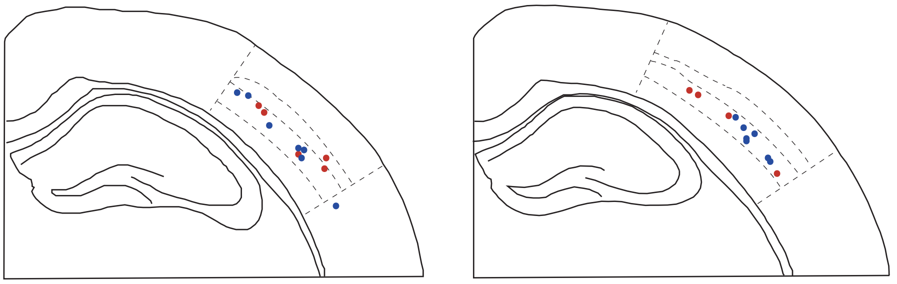

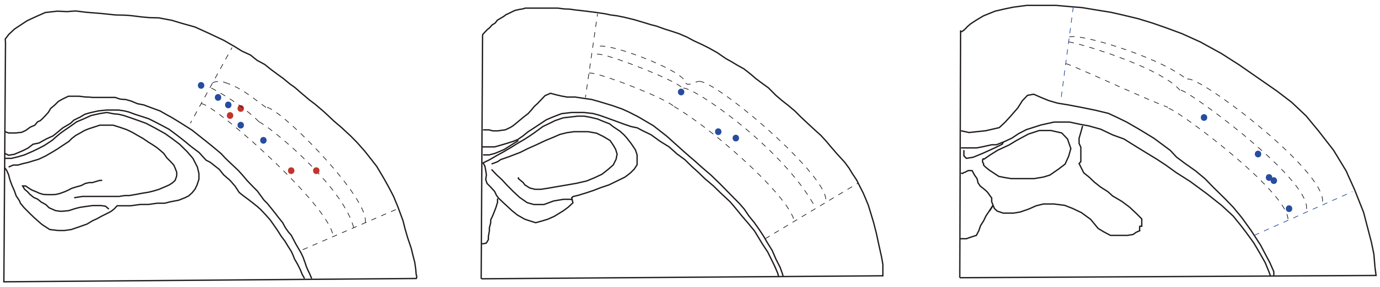

### FIgure S11

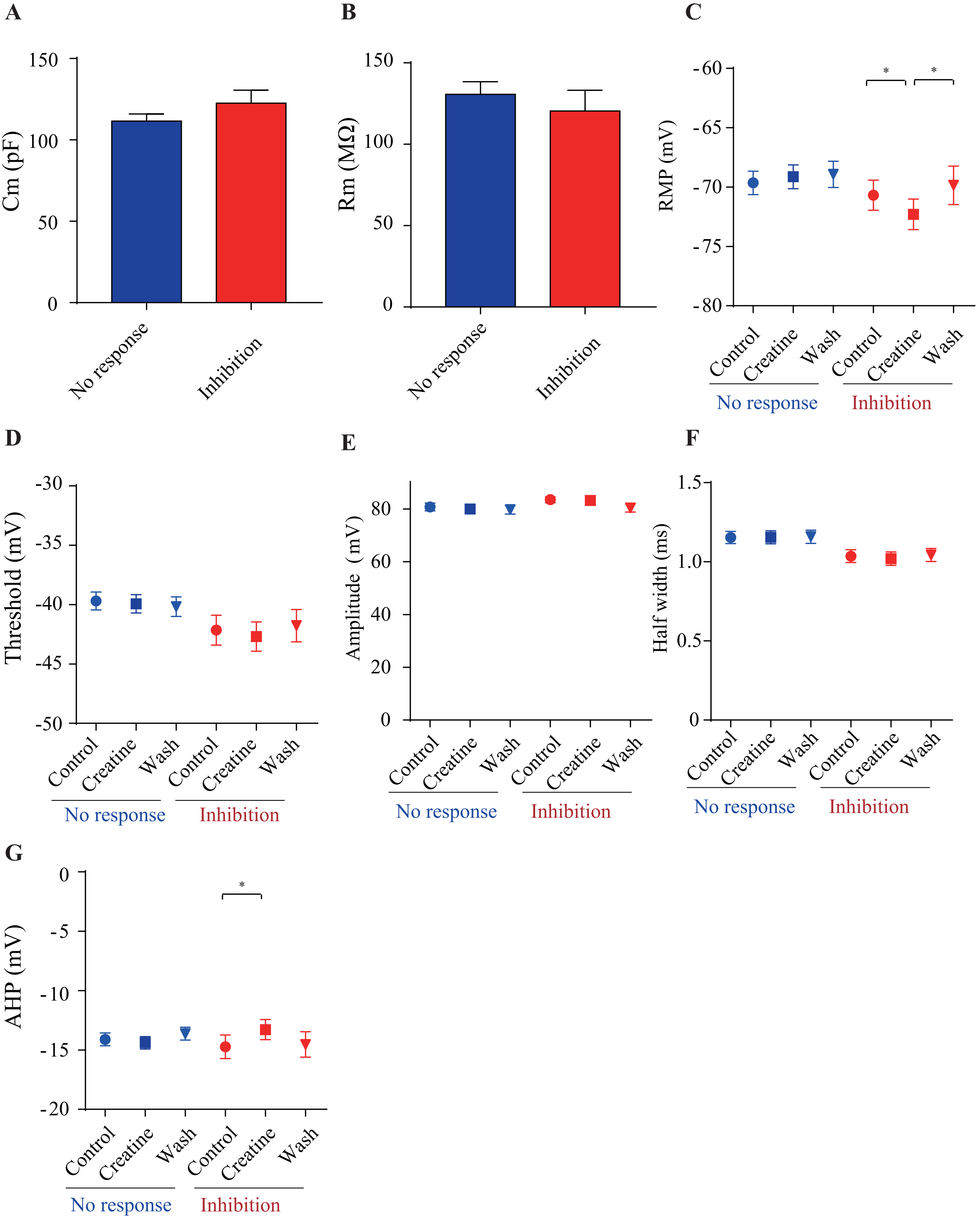

### FIgure S12

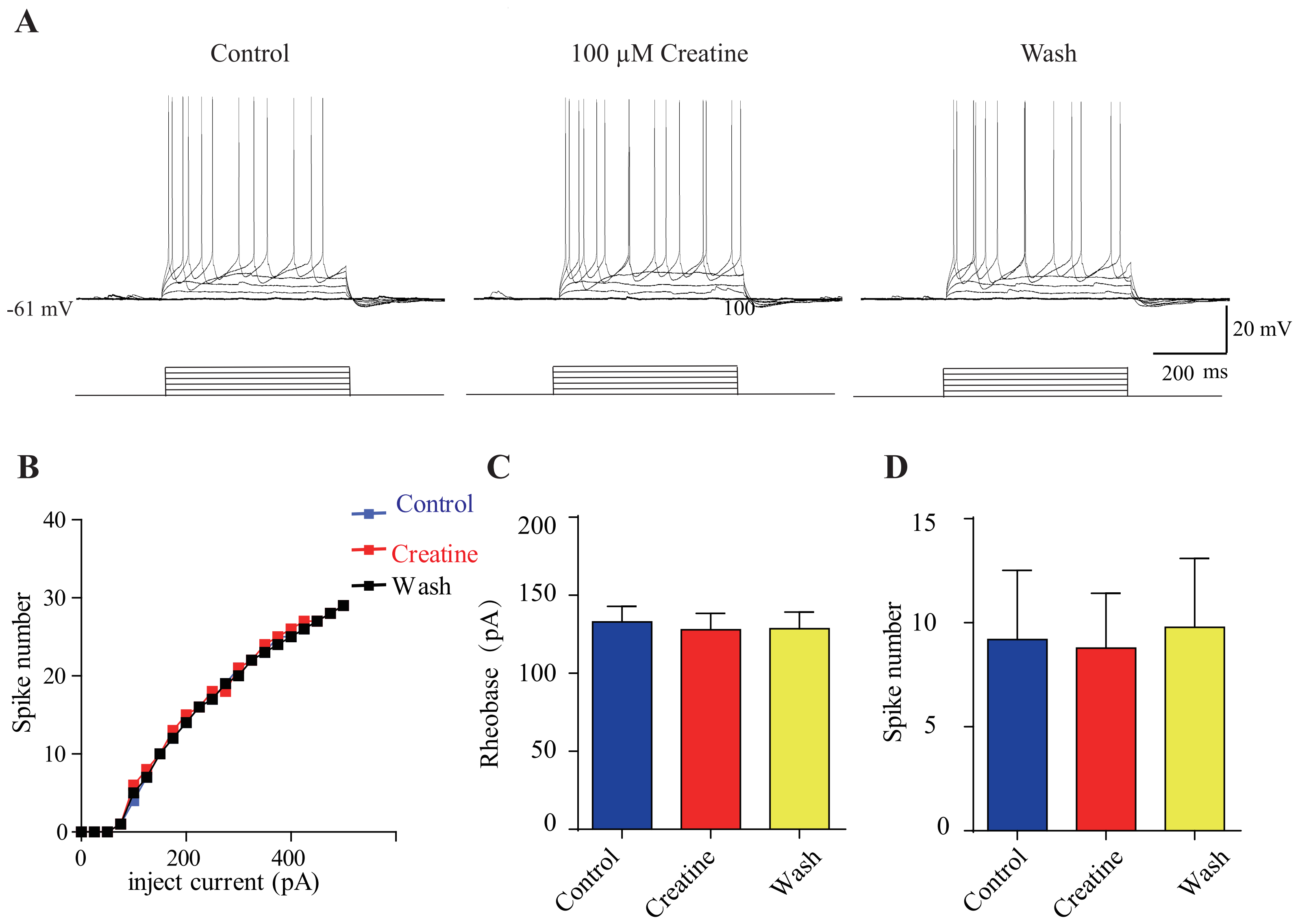
